# Astrocytic urea cycle detoxifies Aβ-derived ammonia while impairing memory in Alzheimer’s disease

**DOI:** 10.1101/2021.10.15.464517

**Authors:** Yeon Ha Ju, Mridula Bhalla, Seung Jae Hyeon, Ju Eun Oh, Seonguk Yoo, Uikyu Chae, Jae Kwon, Wuhyun Koh, Jiwoon Lim, Yongmin Mason Park, Junghee Lee, Il-Joo Cho, Hyunbeom Lee, Hoon Ryu, C. Justin Lee

## Abstract

Alzheimer’s disease (AD) is one of the foremost neurodegenerative diseases, characterized by beta-amyloid (Aβ) plaques and significant progressive memory loss. In AD, astrocytes are known to take up and clear Aβ plaques. However, how Aβ induces pathogenesis and memory impairment in AD remains elusive. We report that normal astrocytes show non-cyclic urea metabolism, whereas Aβ-treated astrocytes show switched-on urea cycle with upregulated enzymes and accumulated entering-metabolite aspartate, starting-substrate ammonia, end-product urea, and side-product putrescine. Gene-silencing of astrocytic ornithine decarboxylase-1 (ODC1), facilitating ornithine-to-putrescine conversion, boosts urea cycle and eliminates aberrant putrescine and its toxic by-products ammonia, H_2_O_2_, and GABA to recover from reactive astrogliosis and memory impairment in AD model. Our findings implicate that astrocytic urea cycle exerts opposing roles of beneficial Aβ detoxification and detrimental memory impairment in AD. We propose ODC1-inhibition as a promising therapeutic strategy for AD to facilitate removal of toxic molecules and prevent memory loss.

## Introduction

Alzheimer’s disease (AD) is a progressive neurodegenerative disease that gradually destroys one’s memory, cognition and even life (2021). The main feature of AD is the accumulation of abnormal protein aggregates, such as beta-amyloid (Aβ) and tau tangles, resulting in neuronal death (Taylor et al., 2002). Other pathological features of AD are the morphological changes of glial cells, especially astrocytes, which usually become hypertrophied and reactive (Anderson et al., 2014; Chun and Lee, 2018). Astrocytic reactivity is a pathological hallmark of astrocytes in response to injury or toxic molecules (Escartin et al., 2021). Reactive astrocytes are early-onset markers of pre-symptomatic AD, appearing before the onset of Aβ- and tau-induced neuronal dysfunction (Leclerc and Abulrob, 2013). In this regard, there is an urgent need for in-depth investigation of the reactive astrocytes that represent the initial alteration in AD.

Reactive astrocytes manifest not only morphological changes but also various transcriptional, metabolic, and functional changes (Chun and Lee, 2018; Escartin et al., 2021). In AD mouse model, reactive astrocytes are shown to display metabolic and functional changes to produce aberrant levels of the neurotransmitter γ-aminobutyric acid (GABA) by monoamine oxidase B (MAO-B) from putrescine via the putrescine-degradation pathway, leading to memory impairment (Jo et al., 2014; Park et al., 2019). Furthermore, toxic by-products such as ammonia and H_2_O_2_ are produced by MAO-B in this pathway, leading to neurodegeneration in AD (Chun et al., 2020). However, there remains the imperative question of how putrescine accumulates from Aβ in reactive astrocytes.

Putrescine, the major precursor of GABA in reactive astrocytes, can be synthesized from two precursor metabolites (Wunderlichova et al., 2014): arginine and ornithine (Morris, 2002; Takiguchi and Mori, 1995). Both metabolites are essential components of the urea cycle (Cohen, 1981; Krebs, 1982). Urea cycle, originally identified in the liver, is one of the principal metabolic pathways that convert the highly toxic ammonia to less toxic urea (Cohen, 1981; Meijer et al., 1990; Morris, 2002). Interestingly, the human brain also contains a comparable concentration of urea (Buniatian and Davtian, 1966). Even more strikingly, an excessive level of toxic ammonia and urea are detected in the brain of AD patients (Adlimoghaddam et al., 2016; Xu et al., 2016). However, the presence of urea cycle in the brain and its pathophysiological effects on memory impairment in AD has not been fully investigated yet.

In this study, we hypothesized that Aβ induces upregulation of the astrocytic urea cycle to detoxify excessive substrate ammonia, producing end-product urea, aberrant side-product putrescine and its by-products ammonia, H_2_O_2_ and GABA, resulting in memory impairment in AD. To investigate this hypothesis, we first examined RNA-sequencing (RNASeq) to detect the changes of gene expression levels in Aβ-treated astrocytes, as well as in the human AD patient brains. We directly measured the ammonia and urea concentrations *in vitro* and developed a novel urea sensor to analyze real-time urea concentration in AD mouse model brain. We performed cell-type specific gene-silencing of various key enzymes in AD culture and animal models to assess urea cycle-related and putrescine-degradation pathway-related metabolites. We also measured GABA production and release and its effects on synaptic transmission by electrophysiology and further examined the behavioral effects of gene-silencing on memory in AD mouse model. Indeed, we demonstrate the existence of urea cycle in reactive astrocytes of AD and the beneficial effect of inhibiting putrescine production on memory impairment in AD.

## Results

### Urea cycle enzymes are upregulated in AD-like conditions

To investigate the existence of urea cycle in astrocytes, we firstly examined the urea cycle pathway-related genes. The conventional urea cycle consists of 5 enzymatic steps, starting from carbamoyl phosphate synthetase1 (CPS1), ornithine transcarbamylase (OTC), argininosuccinate synthetase1 (ASS1), argininosuccinate lyase (ASL) and arginase 1 (ARG1) (Fig. 1A). Overall, the end-product from the ARG1 reaction is ornithine, which can be further converted to putrescine by ornithine decarboxylase 1 (ODC1) or cycle back to citrulline (Fig. 1A). We screened the mRNA level of each enzyme by qRT-PCR in primary astrocyte cultures (Fig. 1B). The basal mRNA level of each enzyme was normalized to *Gapdh* expression level. We found that mRNA for all urea cycle-related enzymes were detected by qRT-PCR with CT value under 34. However, we observed that *Cps1, Otc, Adc* and *Agmat* showed very low expression in normal astrocytes, indicating that the astrocytic urea cycle exhibited different non-cyclic characteristics under normal condition compared to conventional urea cycle pathway. Because *Adc1* and *Agmat* showed very low expression in astrocytes (Fig. 1B), we concluded that the ARG1-to-ODC1 pathway could be the major pathway for putrescine production. To investigate the change in urea cycle-related mechanisms by Aβ treatment, we assessed the Next Generation RNASeq of control and 5-day Aβ treated astrocytes (Fig. 1C). RNASeq with pathway analysis revealed that urea cycle and urea metabolic process were upregulated upon Aβ treatment (Fig. 1D). In addition, we observed that autophagy-related genes (*Atg5, Atg7, Atg12, Lc3A/B, Sqstm1*) and urea cycle-related genes (*Ass1, Asl, Odc1*) were increased by Aβ treatment through differential expression analysis, whereas *Oat* was decreased, indicating that conversion of ornithine to putrescine was preferred over glutamate (Fig. 1E).

**Figure 1.**
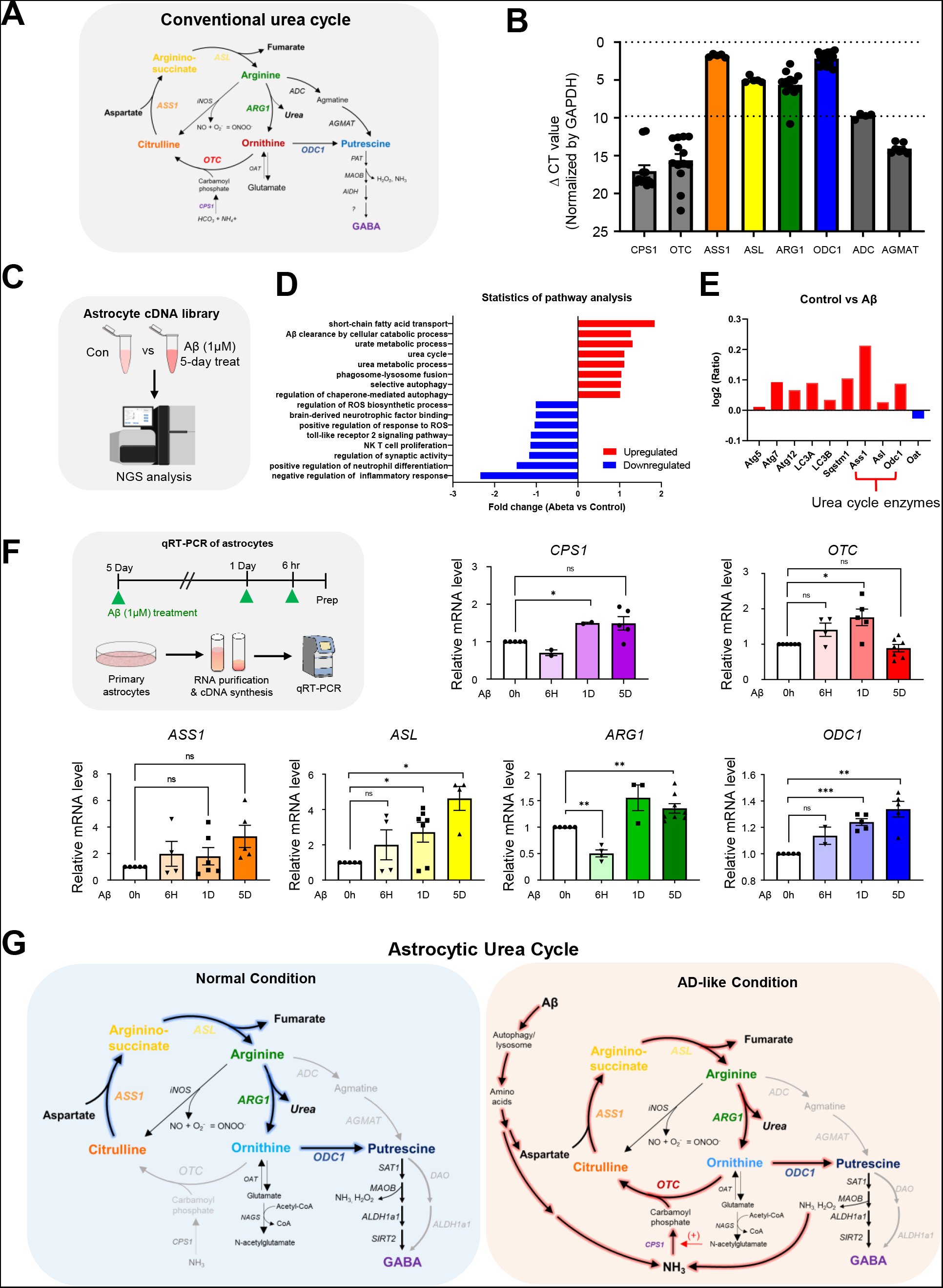
Urea cycle enzymes are upregulated in AD conditions. **(A)** Schematic diagram of the conventional urea cycle. **(B)** Basal mRNA level normalized to GAPDH. **(C)** Experimental diagram of RNASeq. **(D)** Pathway analysis of RNASeq data from Aβ treated astrocytes (1μM, 5days) compared with control astrocytes. **(E)** Differential expression analysis of genes using RNASeq in astrocytes upon Aβ treatment. **(F)** Left, Experimental timeline for qRT-PCR after 5-day, 1-day and 6-hour treatment of Aβ compared to control astrocytes. Right, Bar graphs for relative mRNA level of urea cycle enzymes (*Cps1*, *Otc*, *Ass1*, *Asl*, *Arg1* and *Odc1*), normalized to control. **(G)** Schematic diagram of astrocytic urea cycle (Left, normal condition/ Right, AD-like condition). Data represents Mean ± SEM. *, p<0.05; **, p<0.01; ***, p<0.001.

To verify the obtained RNASeq results, we examined the qRT-PCR of Aβ-treated astrocytes. In this case, the analysis was performed after 4 different durations of Aβ treatment conditions: 0-hour, 6-hour, 1-day and 5-day treatments. As a result, all urea cycle-related genes such as, *Cps1, Otc*, *Ass1, Asl, Arg1* and *Odc1* showed increased tendency with 5-day Aβ treatment (Fig. 1F), indicating that astrocytic urea cycle behaves like the conventional urea cycle under AD-like condition. Taken together, these results indicate that, under normal condition, astrocytes have a unique non-cyclic urea metabolic pathway that lacks CPS1 and OTC enzymes with disrupted cycling, whereas under AD-like condition, the astrocytic urea metabolism reverts to the conventional urea cycle with the expression of most urea cycle-related genes elevated and the production of putrescine from ornithine (Fig. 1G).

### Astrocytic OTC, ARG1 and ODC1 are increased in the hippocampus of AD patients

To assess the clinical importance of the urea cycle in AD patients, we obtained transcriptome data from post-mortem brain samples of 8-10 individuals with AD and control human subjects (Fig. 2A, B). Using KEGG pathway analysis, we found that amino acids biosynthesis and urea cycle-related pathways are upregulated in AD patients (Fig. 2A). Furthermore, we observed the increased level of *OTC, ARG1* and *ODC1* mRNA expression in AD patients by transcriptome analysis. We found that *OTC, ARG1* and *ODC1* transcriptome exhibited the similar upregulated patterns in primary astrocyte cultures and AD patients. We observed that the level of *OTC* was about 100 times lower than *ODC1*, consistent with astrocyte cultures (Fig. 1B). Next, to examine protein levels of OTC, ARG1 and ODC, we performed immunohistochemistry with post-mortem hippocampal tissues of AD patients and normal subjects (Fig. 2C–H). We found an increase of GFAP-positive reactive astrocytes in AD-patient brains compared to the normal group (Fig. 2C, E, G). Furthermore, we found that the expression of OTC, ARG1 and ODC1 was significantly increased and co-localized in GFAP-positive area with increased level in AD brain (Fig. 2D, F, H), indicating that upregulated astrocytic OTC, ARG1 and ODC1 can be used as a pathogenic marker for AD. Because inhibition of OTC is expected to eventually increase, rather than decrease the putrescine concentration, we decided to focus on ARG1 and ODC1, which are the last two steps upstream of putrescine production. Taken together, we propose that in AD conditions, the astrocytic urea cycle shows upregulated ARG1 and ODC1 enzymes, a process that leads to the synthesis of putrescine and eventually GABA.

**Figure 2.**
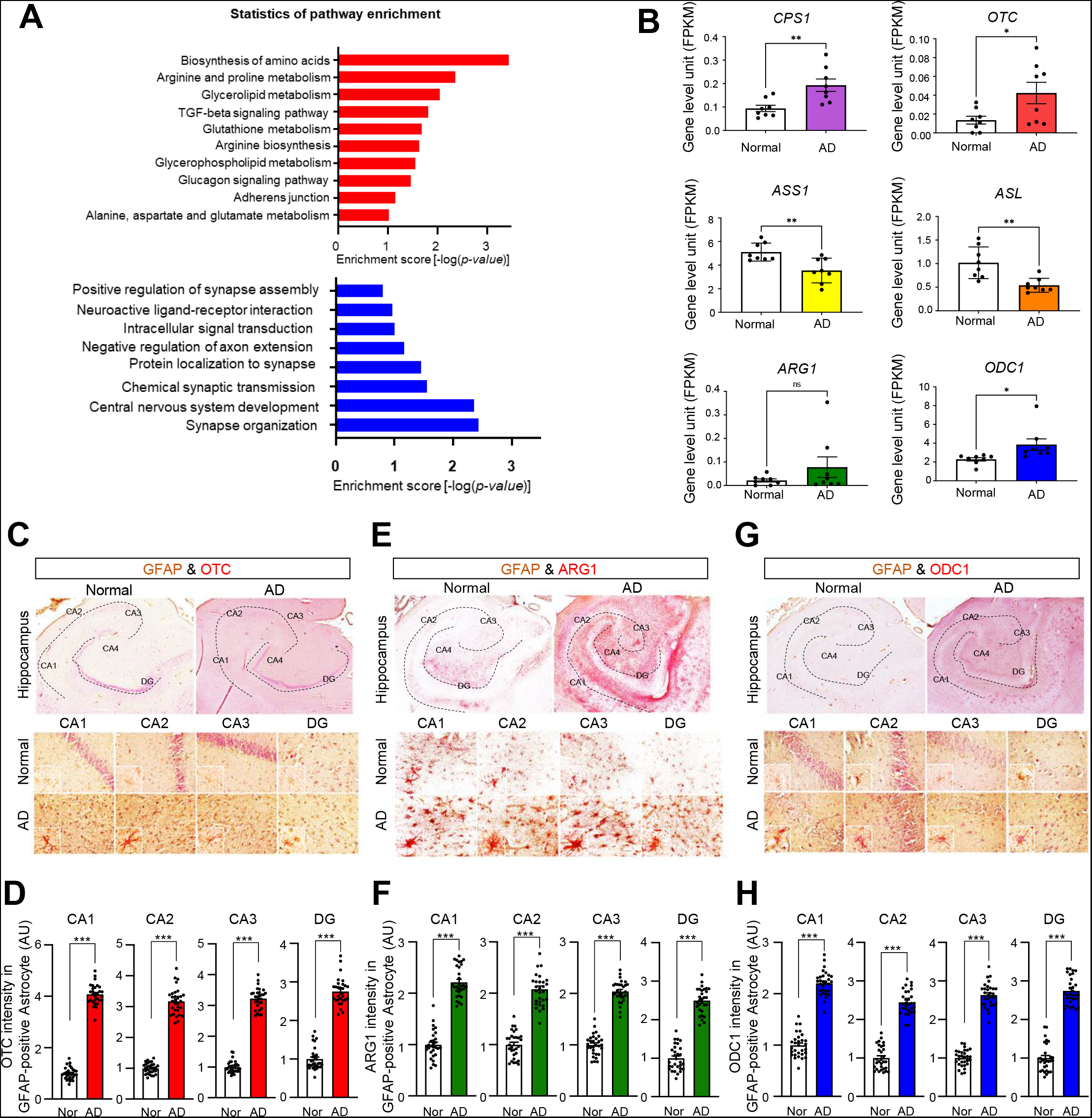
Urea cycle-related genes are elevated in astrocytes of the hippocampus in AD patients. (A) Comparative pathway enrichment analysis of amino acid metabolism- and urea cycle-associated KEGG pathways of in human AD patients.The number of up-(red, top) or down-regulated (blue, bottom) transcriptome signatures involved in each KEGG pathway is shown in the side bar graph. **(B)** Bar graphs for comparison of gene expression units (FPKM) of urea cycle enzymes (*Cps1*, *Otc*, *Ass1*, *Asl*, *Arg1* and *Odc1*). **(C, D)** Representative images (C) and bar graphs (D) of OTC immunoreactivity in GFAP-positive astrocytes of the hippocampus in normal subjects and AD patients. **(E, F)** Representative images (E) and bar graphs (F) of ARG1 immunoreactivity in GFAP-positive astrocytes of the hippocampus in normal subjects and AD patients. **(G, H)** Representative images (G) and bar graphs (H) of ODC1 immunoreactivity in GFAP-positive astrocytes of the hippocampus in normal subjects and AD patients. Data represents Mean ± SEM. *, p<0.05; **, p<0.01; ***, p<0.001.

### Astrocytic urea cycle metabolites, including entering-metabolite, starting-substrate, end-product, side-product, and by-products are upregulated in AD-like conditions

To investigate the direct metabolic changes of astrocytic urea cycle in AD-like conditions, we analyzed the intracellular astrocytic urea cycle metabolites by liquid chromatography/mass spectrometry (LC/MS) analysis in cultured astrocytes with 1μM Aβ treatment (Fig. 3A). We found that aspartate, one of the substrate amine donors in urea cycle (Fig. 1G), was significantly increased by Aβ treatment. Furthermore, the concentration of side-product putrescine and its by-product GABA were significantly increased by Aβ treatment in astrocytes (Fig. 3A), consistent with our previous report (Jo et al., 2014), whereas glutamate, a potential side-product of ornithine (Fig. 1G), showed no change (Fig. 3A). We discovered that the intermediate metabolites of urea cycle, such as citrulline, arginine and ornithine did not change significantly (Fig. 3A), suggesting that these urea cycle metabolites are in a constant flux. In contrast, only aspartate showed significant increase, suggesting that aspartate is the major entering metabolite to the urea cycle from Aβ degradation. We further confirmed ammonia was the starting-substrate for the urea cycle by treating primary cultured astrocytes with ^15^NH_4_Cl (heavy nitrogen ammonium chloride, 1mM) and found that ^15^N-atoms incorporated into ornithine, putrescine and GABA (Fig. S2).

**Figure 3.**
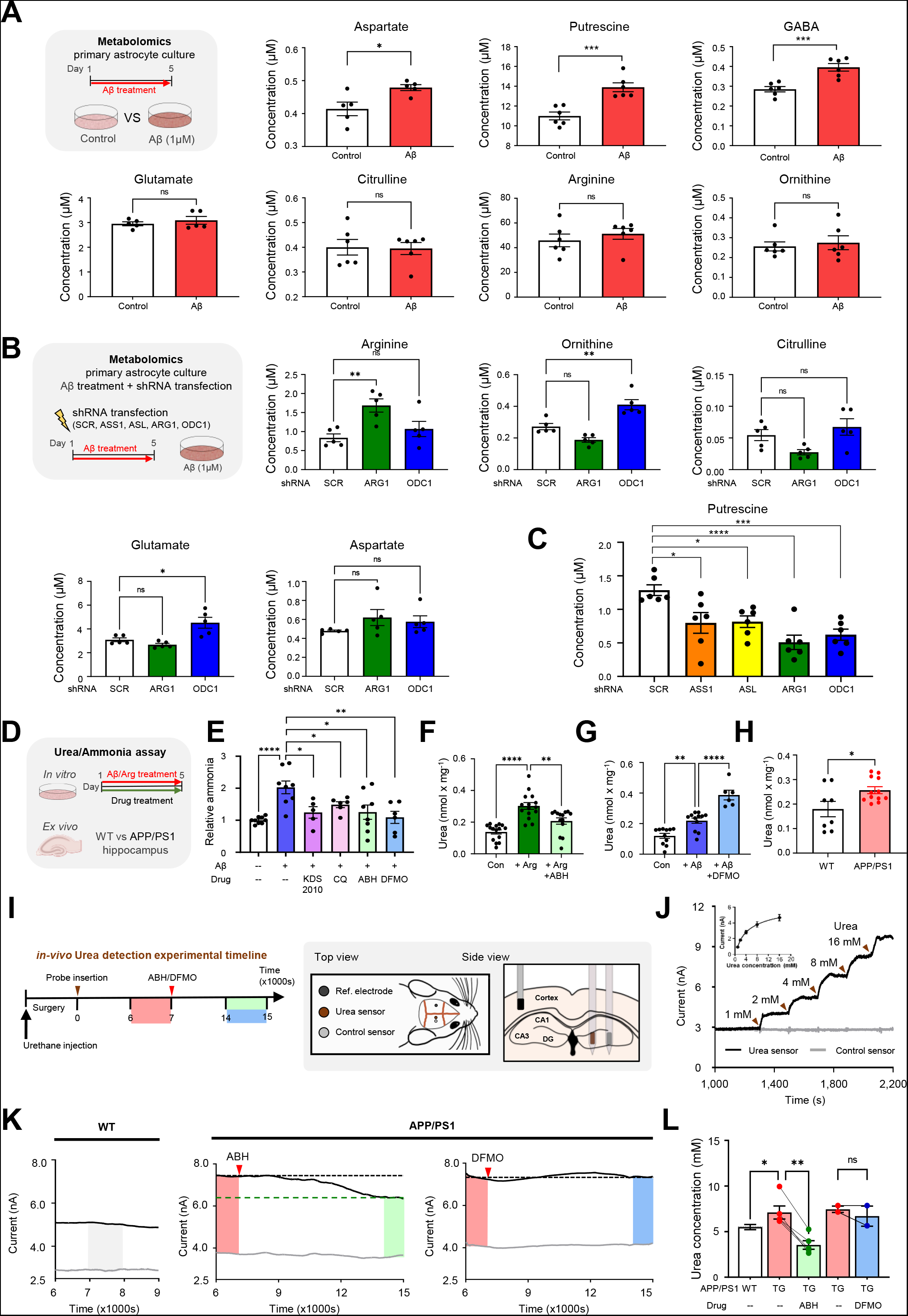
Urea cycle metabolites and urea production are upregulated in AD-like conditions. **(A)** Left, Schematic of metabolite analysis timeline. Right, Bar graphs for concentration of each metabolite from control and Aβ-treated astrocytes (Aspartate, Putrescine, GABA, Citrulline, Arginine and Ornithine; see also Fig. S2). **(B)** Left, Schematic of metabolite analysis timeline. Right, Bar graphs for concentration of metabolites in Scr-, Arg1- and Odc1-shRNA transfected astrocytes. **(C)** Putrescine concentration in Scr-, Ass1-, Asl-, Arg1- and Odc1-shRNA transfected astrocytes (See also Fig. S1). **(D)** Experimental timeline for urea assay and ammonia assay *in vitro* and *ex vivo*. **(E**) Concentration of ammonia in primary cultured astrocytes with Aβ in the presence or absence of CQ, ABH hydrochloride, DFMO or KDS2010. **(F)** Concentration of urea in primary astrocytes with arginine and arginine + ABH hydrochloride treatment. **(G)** Concentration of urea in primary astrocytes with Aβ and Aβ + DFMO treatment. **(H)** Concentration of urea in WT and APP/PS1 mice hippocampal sections, individual dots refer to single brain slice, WT N=4, APP/PS1 N=6. **(I**) Left, Schematic representation of experimental timeline of *in vivo* urea detection experiments in APP/PS1 and WT mice. Right, Representative position of electrode insertion in mouse brain (See also Fig. S3). **(J)** Current response of urea and control sensors upon increasing concentration of urea from 1 mM to 16 mM *in vitro* aCSF solution, brown arrowheads indicate time of application of urea solution having corresponding concentration. Inset, calibration curve for urea sensing showing the dynamic up to 16mM. Mean ± SEM of 11 sensors. **(K)** Representative traces of current recording from urea sensor in WT (Left) and APP/PS1 mouse brain before and after injection of ABH (Middle) or DFMO (Right), red arrowheads indicate time of ABH/DFMO injection, shaded boxes indicate current value used for calculation of urea concentration. **(L)** Bar graph indicating urea concentration as calculated from above traces in APP/PS1 mouse brains, individual dots refer to one animal, connecting lines indicate paired recordings, WT N=4, APP/PS1-ABH N=4, APP/PS1-DFMO N=2. Data represents Mean ± SEM. Individual dots refer to single batch of cultured cells unless otherwise clarified. *, p<0.05; **, p<0.01; ***, p<0.001.

To investigate the flux of metabolites in astrocytic urea cycle, we developed shRNAs for each urea cycle enzyme, including *Ass1*, *Asl*, *Arg1* and *Odc1* (Fig. S1). We first transfected Aβ-treated astrocytes with Arg1- or Odc1-shRNA to inhibit each enzyme by gene-silencing and compare with control Scrambled (Scr) shRNA (Fig. 3B). We found that concentration of arginine was significantly higher in Arg1-shRNA condition (Fig. 3B), suggesting that silencing *Arg1* induces accumulation of arginine, caused by constant flux of astrocytic urea cycle by Aβ treatment (Fig. 3B). Furthermore, ornithine concentration was significantly higher in Odc1-shRNA condition, also confirming the constant flux of astrocytic urea cycle by Aβ treatment. In addition, we observed that concentration of citrulline tended to increase by Odc1-shRNA (Fig. 3B), suggesting that the accumulated ornithine by *Odc1* gene-silencing is circulated back to citrulline. Since aspartate is upstream to ARG1 and ODC1, the concentration of aspartate was not changed upon gene-silencing (Fig. 3B). We also found the increased concentration of glutamate by silencing *Odc1*, raising a possibility that the accumulated ornithine might be veered off to glutamate through OAT pathway. Taken together, these results indicate that astrocytic urea cycle is circulating in a constant flux under AD-like condition.

To investigate whether astrocytic urea cycle is the major upstream source of putrescine production in AD-like condition, we transfected Aβ-treated astrocytes with shRNAs (*Ass1*, *Asl*, *Arg1* or *Odc1*) and examined the putrescine concentration (Fig. 3C). We observed that putrescine concentration was significantly reduced in all conditions, indicating that the astrocytic urea cycle enzymes are involved in putrescine production. Moreover, gene-silencing of *Arg1* and *Odc1* showed most dramatic reduction of putrescine production, indicating that these enzymes are the key modulators of this process. Altogether, these results provide direct evidence that Aβ-treatment increases level of aspartate, the entering metabolite, and putrescine, the end-product of the astrocytic urea cycle, with intermediate metabolites in constant flux, possibly at an elevated rate.

Next, to investigate the source and fate of the substrate ammonia, we performed ammonia assay in AD-like condition (Fig. 3D, E). The possible sources of toxic ammonia entering the urea cycle include elevated amino acid metabolism due to increased protein degradation via Aβ autophagy/lysosome pathway (Ries, 2016) and MAO-B-dependent ammonia production via the putrescine degradation pathway (Jo et al., 2014) (Fig 1G). To dissect the contribution of autophagy/lysosome and MAO-B-dependent pathway, we performed the ammonia assay on Aβ-treated astrocytes with autophagy inhibitor CQ (Chloroquinone, 20uM) (Mauthe, 2018) or recently developed reversible MAO-B inhibitor KDS2010 (100nM) (Park et al., 2019). We observed that MAO-B inhibition drastically reduced the levels of accumulated ammonia while autophagy inhibition partially decreased ammonia levels as compared to Aβ-treatment (Fig. 3E), indicating that the MAO-B-mediated putrescine degradation pathway was the major source of toxic ammonia entering the urea cycle. We then sought to examine the effect of pharmacological inhibition of ARG1 and ODC1 on the accumulation of toxic ammonia in Aβ-treated astrocytes. We performed the ammonia assay on Aβ-treated astrocytes along with ARG1 inhibitor ABH (2(S)-amino-6-boronohexanoic acid - hydrochloride, 10 μM) (Baggio, 1999) and ODC1 inhibitor DFMO (Diflouromethylornithine, 50uM) (LoGiudice, 2018). We found that both ABH and DFMO were able to significantly reduce the accumulation of ammonia from Aβ treatment (Fig. 3E). Taken together, these results suggest the presence of a toxic positive feedback loop, and that inhibition of ARG1, ODC1 or MAO-B can prevent the accumulation of toxic ammonia and limit its re-entrance into the urea cycle.

To investigate whether astrocytic urea cycle produces end-product urea on Aβ treatment, we examined the urea concentration *in vitro* and *ex vivo* through urea assay (Fig. 3D). We measured the cellular content of urea in primary astrocyte cultures on 5-day treatment with arginine (10mM) and ABH (10μM). The cellular content of urea was found to be significantly increased by arginine treatment, which was blocked by ARG1 inhibition (Fig. 3F), indicating that both arginine and ARG1 are key players in urea production. Next, to test the effect of Aβ and ODC1 on urea production, we treated astrocytes with Aβ (1μM) and DFMO (50μM) (Fig. 3G). We found that urea concentration was increased by Aβ treatment, which was further enhanced by DFMO treatment (Fig. 3G), indicating that ODC1 inhibition accelerates the detoxification of ammonia to urea by the urea cycle. These results indicate that ARG1 inhibition disrupts Aβ and ammonia detoxification by the urea cycle whereas ODC1 inhibition enhances it. Taken together, astrocytes show upregulated flux of astrocytic urea cycle and production of end-product urea on Aβ treatment

To further examine the *in vivo* significance of the urea cycle enzymes in AD mouse model, we developed a novel urea sensor probe which allowed for real-time measurement of urea in the brain of anesthetized WT and transgenic APP/PS1 (TG) mice, an animal model of AD (Fig S3, Fig 3I). Before using the novel urea sensor probe, we firstly measured the intracellular content of urea in WT and APP/PS1 mice hippocampal tissue (Fig. 3H) using an enzyme-based urea assay. Consistent with a previous report (Xu et al., 2016), we observed that urea concentration was significantly higher in APP/PS1 mice than WT mice (Fig 3H), confirming that the urea cycle is indeed upregulated in AD mice. We then employed the newly developed urea sensor to measure urease-induced current (Figure S3) in the dentate gyrus (DG) of WT and APP/PS1 mice and calculated extracellular urea concentration based on pre-calibration of the sensor *in vitro* (Fig. 3J). We observed that APP/PS1 mice had a higher urea concentration than WT mice (Fig. 3K and L). Current was then measured in real-time before and after injection of ABH (10mg per kg animal weight) or DFMO (600mg per kg animal weight) in APP/PS1 mice in paired experiments (Fig. 3I). After i.p injection of ABH, we found that urea-induced current gradually and significantly reduced within about 2 hours (Fig 3K), translating as reduced extracellular urea concentration in the brain (Fig 3L). Injection of DFMO, however, induced no such change (Fig. 3K and L). These results indicate ARG1 is a key enzyme in the production of urea from detoxification of Aβ in AD mouse brain.

### ARG1 and ODC1 are essential upstream enzymes for by-product GABA

To test whether urea cycle is directly involved in production of GABA by putrescine degradation, we developed and utilized shRNAs for urea cycle enzymes to silence urea cycle genes in primary astrocyte cultures (Fig. S1, 4A–C). Urea cycle enzyme shRNAs (*Ass1*, *Asl*, *Arg1* and *Odc1*) were individually transfected in astrocytes treated for 5-days with Aβ and stained for GFAP and GABA by immunocytochemistry (ICC). As a result, GABA was significantly decreased by gene-silencing the enzymes involved in urea cycle (Fig. 4B and C), indicating that urea cycle is upstream to the GABA synthesis pathway.

**Figure 4.**
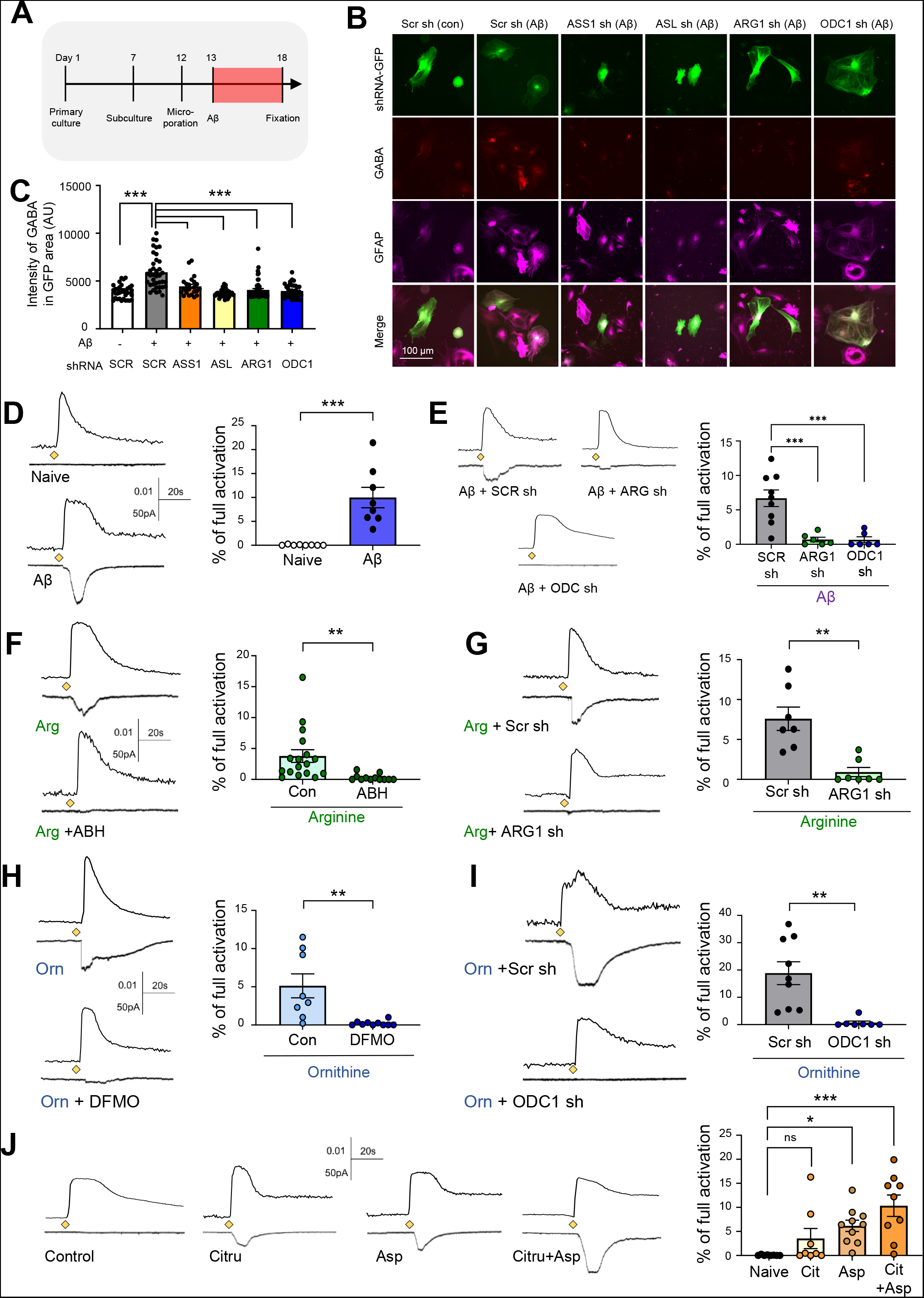
ARG1 and ODC1 are major enzymes for GABA production. **(A)** Experimental timeline of immunocytochemistry. **(B)** Immunostaining of GFAP and GABA in shRNA-transfected astrocytes (Scr-, Ass1-, Asl-, Arg1- and Odc1-shRNA conditions; see also Fig S1). **(C)** Quantification of GABA in GFAP positive area (AU). (D-J) Representative traces and bar graphs of sensor current induced by GABA from primary cultured astrocytes in different conditions. (D) Astrocytes with or without Aβ-treatment. (E) Aβ-treated astrocytes transfected with Scr-, Arg1- or Odc1-shRNA. **(F, G)** Arginine-treated astrocytes in the presence or absence of ABH hydrochloride (F) and transfected with Scr- or ARG1-shRNA (G). **(H, I)** Ornithine-treated astrocytes in the presence or absence of DFMO (H) and transfected with Scr- or ODC1-shRNA. **(J)** Naive, Citrulline-, Aspartate- and citrulline + Aspartate-treated astrocytes (See also, Fig S4) Diamonds indicate poking of the astrocytes. Data represents Mean ± SEM. Individual dots refer to cells unless otherwise clarified. **, p < 0.01 (Student’s t-test); ***, p < 0.001 (One-way analysis of variance (ANOVA) and Sheffe’s test).

To investigate whether ARG1 and ODC1 are key modulator enzymes for producing GABA under AD-like condition, we performed sniffer-patch experiments from Aβ-treated cultured astrocytes to examine the immediate release of GABA in real-time from a solitary astrocyte (Jo et al., 2014). On poking an astrocyte, increased calcium signal by TRPA1 channel is measured (Oh, 2019) and released GABA can be measured as current recorded from GABAc receptor-expressing HEK cells. Inward GABA current was normalized to the full activation current by 100μM GABA treatment (Fig. S4A and B). After 5-day treatment of Aβ, astrocytes showed a significant release of GABA (Fig. 4D), as previously described (Jo et al., 2014), which was almost completely eliminated by gene-silencing of *Arg1* and *Odc1* using shRNA (Fig. 4E). We then incubated primary cultured astrocytes with arginine (10mM), the substrate for ARG1, and compared the control with ABH treatment (10μM). We found that arginine treatment increased GABA release, which was blocked by ARG1 inhibitor (Fig. 4F). Consistently, GABA release due to arginine incubation was significantly decreased by gene silencing of *Arg1* (Fig. 4G), indicating that both pharmacological and genetic inhibition of ARG1 blocks the arginine-induced production of GABA. Next, when we treated the culture with ornithine (10mM), which is the precursor of putrescine, GABA current was induced, which was significantly eliminated by ODC1 inhibitor, DFMO (50μM, Fig. 4H). Finally, increased GABA current by ornithine was significantly decreased by gene-silencing of *Odc1* (Fig. 4I), indicating that ODC1 is essential for ornithine-induced GABA production. Inhibition of autophagy by CQ (20μM) also blocked Aβ-induced GABA production (Fig. S4A–C), further confirming the upstream role of autophagy in putrescine, and consequently GABA production. Additionally, results from H_2_O_2_ assay further supported that ODC1 and ARG1 inhibition reduced Aβ-induced H_2_O_2_ production and oxidative stress (Fig. S4D).

Since citrulline and aspartate are also essential to initiate the urea cycle, we incubated primary cultured astrocytes with citrulline (10mM) and aspartate (10mM) independently and together. We found that citrulline alone had no significant effect on GABA production and release, whereas aspartate significantly increased it (Fig. 4J), indicating that accumulation of entering-metabolite aspartate could increase the flux of the urea cycle, leading to more GABA production. Interestingly, citrulline with aspartate showed the greatest increase in GABA current, suggesting aspartate and citrulline together facilitate the urea cycle. Taken together, these results indicate that increased flux of the urea cycle, which is upstream to the putrescine degradation pathway, causes increased GABA and H_2_O_2_ production in AD-like condition, a process in which enzymes ARG1 and ODC1 are key players.

### Aberrant GABA and putrescine production is reduced by gene-silencing of *Arg1* and *Odc1* in APP/PS1

To examine the ODC1 level in APP/PS1 transgenic mice, we performed immunohistochemistry (IHC) with antibodies against GFAP and ODC1 (Fig. 5A). We found that ODC1 was minimally stained in the astrocytes of WT mice, whereas APP/PS1 mice showed a remarkable appearance of ODC1 in the GFAP-positive area in the hilus of the dentate gyrus (DG) but not in the granule cell (GC) layer (Fig. 5A). Consistent with human data (Fig. 2), these results raise ODC1 as a new marker for AD, due to its high disease-associated expression. To see the effect of Odc1-shRNA *in vivo*, we injected a lentivirus carrying pSicoR-Odc1shRNA-GFP construct into the dentate gyrus (DG) in WT and APP/PS1 mice. Three weeks after virus injection, we sacrificed the animal and performed IHC using antibodies against GFAP and ODC1. We observed that upregulated astrocytic ODC1 was completely reversed by Odc1-shRNA, (Fig. 5B). These results depict the high efficiency of the Odc1-shRNA construct *in vivo* as well as the specificity and reliability of the ODC1 antibody. In contrast, we were unable to obtain a suitable antibody against mouse ARG1.

**Figure 5.**
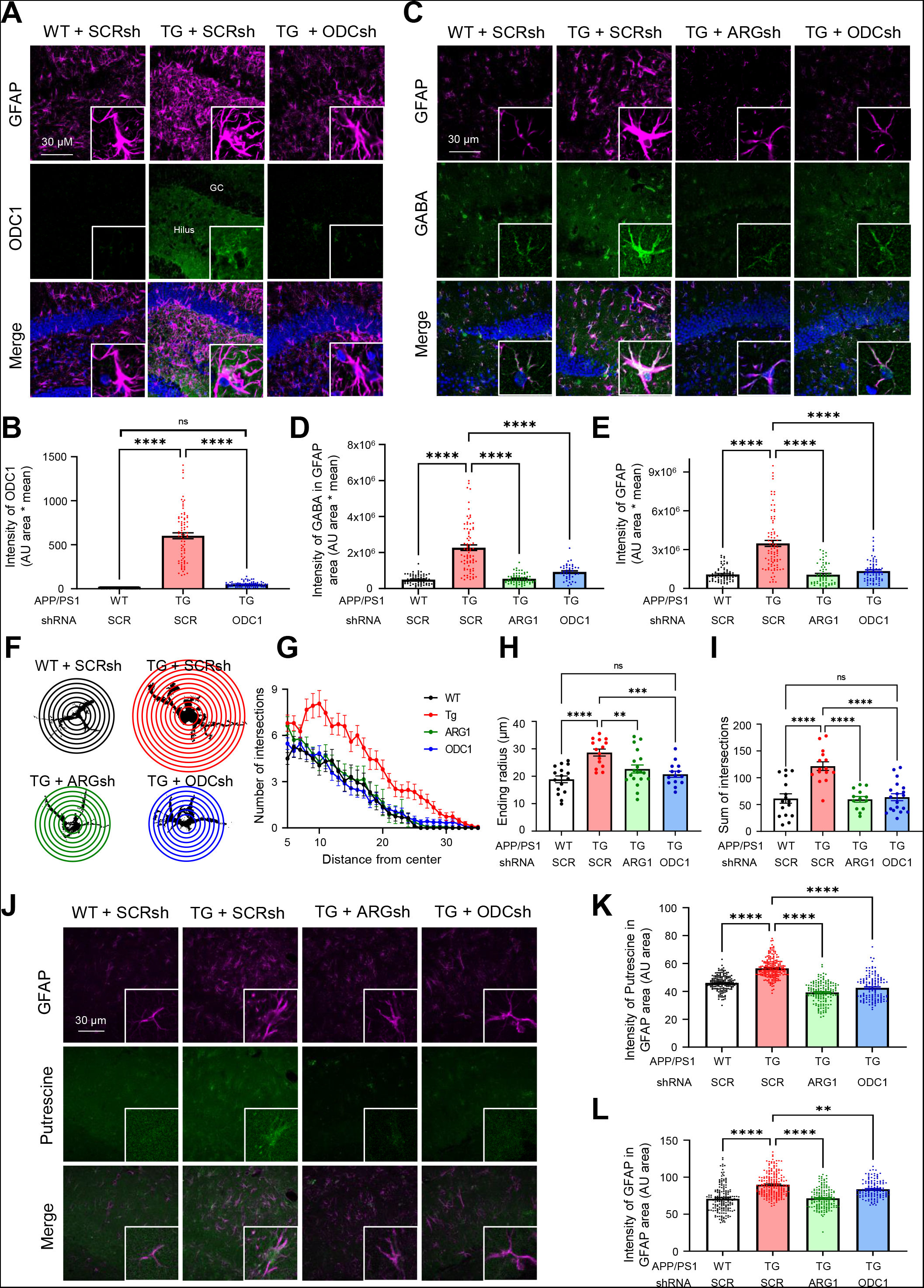
Aberrant GABA production is reduced by inhibition of ARG1 and ODC1. **(A)** Immunostaining for ODC, GFAP and DAPI in Lenti-pSicoR-Scrsh-GFP/ Lenti-pSicoR-ODC1sh-GFP virus injected APP/PS1 mice (pseudo-colored). **(B)** Mean intensities of ODC1 in GFAP-positive area. **(C)** Immunostaining for GFAP, GABA and DAPI in Lenti-pSicoR-Scrsh-GFP/ Lenti-pSicoR-ARG1sh-GFAP/ Lenti-pSicoR-ODC1sh-GFP virus injected WT or APP/PS1 mice (pseudo-colored). Inset: magnified images. **(D, E)** Mean intensities of GABA (D) and GFAP (E) in GFAP-positive areas (AU area * mean value). **(F)** Representative image for Sholl analysis of an astrocyte in the hilus of the DG from the GFAP-stained images in C. Interval of concentric circles is 10μm. **(G)** Average number of intersections in Sholl analysis with respect to the distance from the center of each GFAP-stained cell. **(H, I)** Bar graph showing the ending radius (H) and summarized number of intersects (I) by Sholl analysis of the immunostained GFAP signal. **(J)** Immunostaining for GFAP and Putrescine in AVV-GFAP-Cre-mCh with Lenti-pSico-Scrsh-GFP/Lenti-pSico-ARG1sh-GFP/Lenti-pSico-ODC1sh-GFP virus-injected APP/PS1 mice (pseudo-colored). **(K, L)** Mean intensities of Putrescine (K) and GFAP (L) in GFAP-positive areas. **, p < 0.01; ***, p < 0.001; ****, p < 0.0001, one-way ANOVA with Tukey’s multiple comparisons test. Data represents Mean ± SEM. Numbers and individual dots refer to cells unless otherwise clarified.

Next, to test whether ARG1 and ODC1 are key upstream enzymes leading to GABA synthesis in the animal model of AD, we injected viruses carrying pSicoR-Arg1shRNA-GFP or pSicoR-Odc1shRNA-GFP into WT and APP/PS1 mice (Fig. 5C). We found that APP/PS1 mice showed aberrant GABA production in GFAP-positive astrocytes, which was reverted to WT levels by Arg1- and Odc1-shRNAs (Fig. 5D), indicating that ARG1 and ODC1 are key upstream enzymes in astrocytic GABA synthesis. In addition, upregulated levels of GFAP in APP/PS1 mice were significantly reduced in Arg1- or Odc1-shRNA-injected mice (Fig. 5E). Detailed Sholl analysis of astrocytes (Fig. 5F) revealed that the increased ending radius (Fig. 5H) and sum of intersects (Fig. 5G and I) of the astrocytes from APP/PS1 mice were also rescued by *Arg1* and *Odc1* gene-silencing, indicating reduced astrogliosis. Given that the increased GABA observed in reactive astrocytes is by MAO-B mediated degradation of putrescine (An, 2021; Jo et al., 2014), we further sought to examine the level of side-product putrescine upon astrocyte-specific gene-silencing of *Arg1* and *Odc1* in APP/PS1 mice. We injected lentivirus carrying Cre-dependent pSico-Arg1shRNA-GFP or pSico-Odc1shRNA-GFP along with AAV-GFAP-Cre-mCherry virus into the DG of WT or APP/PS1 mice and performed IHC for putrescine and GFAP (Fig. 5J). We found that putrescine and GFAP levels were significantly increased in the APP/PS1 mice and were recovered back to WT levels on astrocyte-specific gene-silencing of *Arg1* and *Odc1* (Fig. 5K). Taken together, these results indicate that ARG1 and ODC1 are essential for putrescine- and consequently, GABA-synthesis, as well as reactive gliosis in AD. Consistent with data from Aβ-treated primary cultured astrocytes, (Fig. 4), these results implicate that the astrocytic urea cycle is upstream to the GABA synthesis pathway in AD mouse model.

### Suppressing ARG1 and ODC1 enzymes in astrocytes completely restores impaired spike probability and memory deficit in APP/PS1 mice

To examine the level of astrocytic GABA release by pharmacological inhibition of ARG1 and ODC1 enzymes in APP/PS1 mice, we performed whole-cell patch clamp recordings in DG granule cells after incubation with ARG1 inhibitor ABH (1mM) and ODC1 inhibitor DFMO (1mM) for more than 2 hours (Fig. 6A). We observed that both ABH- and DFMO-incubated slices showed significant reduction of tonic GABA level, with no significant difference in sIPSC frequency and sIPSC amplitude (Fig. 6B), providing strong evidence that ARG1 and ODC1 were involved in GABA synthesis in astrocytes but not neurons. Accordingly, to examine the change in tonic GABA current by astrocyte-specific gene-silencing of *Arg1* and *Odc1* in APP/PS1 mice, we injected lentivirus carrying Cre-dependent pSico-Arg1shRNA-GFP or pSico-Odc1shRNA-GFP with AAV-GFAP-cre-mCherry virus into the DG of WT or APP/PS1 mice (Fig. 6C, D). We observed significantly increased tonic GABA current in APP/PS1 mice compared to WT mice, as previously described (Jo et al., 2014). Interestingly, this increased tonic GABA current was significantly reduced to WT levels by astrocyte-specific Arg1- and Odc1-shRNA expression, with no differences in sIPSC amplitude and frequency. These results indicate that pharmacological and genetic inhibition of ARG1 and ODC1 selectively affects astrocytic tonic GABA release, not synaptic release.

**Figure 6.**
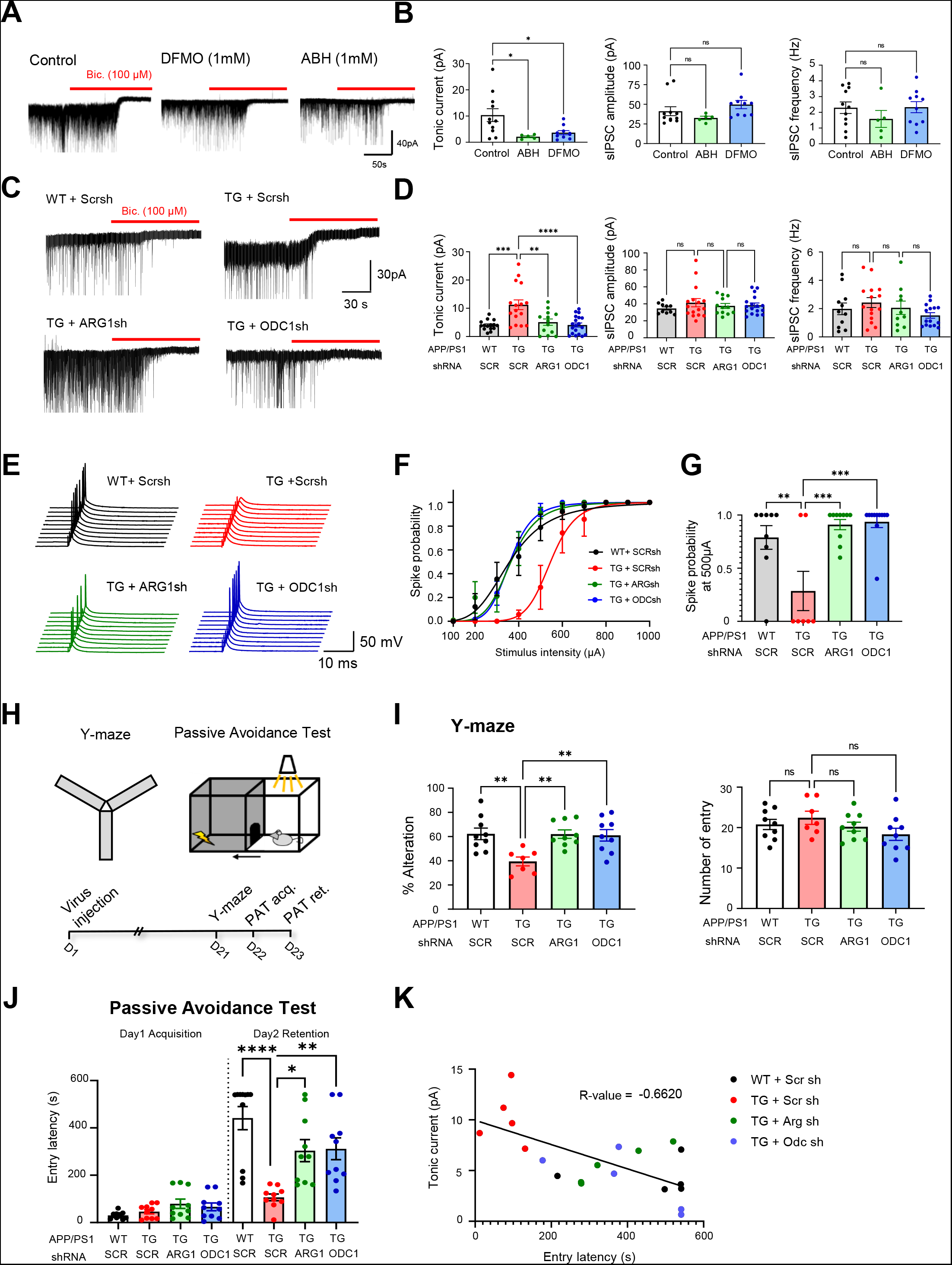
Astrocytic ARG1 and ODC1 are increased in the hippocampus of AD patients. **(A)** Representative trace of GABAA receptor-mediated currents recorded from granule cells in the DG in APP/PS1 brain sections incubated with DFMO (1mM) or ABH (1mM). The red bar indicates application of the GABAA receptor antagonist bicuculline (Bic) (100 μM). **(B)** Comparative bar graphs of tonic GABA current amplitude, sIPSC frequency and amplitude before bicuculline treatment. **(C)** Representative trace of tonic GABA current in WT or APP/PS1 mice injected with AAV-GFAP-Cre-mCh and Lenti-pSico-ScrshRNA/ARG1shRNA/ODC1shRNA-GFP viruses. **(D)** Comparative bar graphs of amplitude of tonic GABA current, frequency and amplitude of sIPSC. **(E)** Representative traces of evoked spikes in hippocampal slices obtained from virus-injected WT and APP/PS1 mice in response to electrical stimulation of the perforant path (0.1 Hz, 100 μs, 100 to 1,000 μA). **(F)** Summary graph of spike probability versus stimulus intensity in Fig. 5E. **(G)** Comparison of spike probability at 500μA stimulation in virus-injected mice. **(H)** Schematic representation of the experimental timeline for behavior test. **(I)** Y-maze test of virus-injected WT and APP/PS1 mice. Bar graphs representing alteration % and number of entries, individual dots represent one animal. **(J)** Passive avoidance test of virus-injected WT and APP/PS1 mice. Bar graphs representing latency to enter dark chamber during passive avoidance test, individual dots represent one animal. **(K)** Correlation analysis between tonic GABA current and Entry latency (memory impairment). **, p < 0.01; ***, p < 0.001; ****, p < 0.0001; one-way ANOVA with Tukey’s multiple comparisons test. Data represents Mean ± SEM. Individual dots refer to cells unless otherwise clarified.

Under AD conditions, tonic GABA release is known to inhibit synaptically elicited action potential firing (Jo et al., 2014). To investigate the role of astrocytic ARG1 and ODC1 in this inhibition, we performed whole-cell current clamp recordings in these virus-injected mice to calculate the spike probability of synaptically elicited action potentials riding on top of EPSPs upon electrical stimulation of perforant pathways as previously described (Jo et al., 2014) (Fig. 6E). Consistent with previous report (Jo et al., 2014), the spike probability was impaired in APP/PS1 mice compared to WT mice (Fig. 6F). Importantly, this impaired spike probability was fully rescued by gene-silencing of astrocytic *Arg1* and *Odc1* (Fig. 6F, G). These results demonstrate that astrocytic ARG1 and ODC1 are necessary for the elevated tonic GABA release and impaired spike probability observed in AD mice.

To further investigate the memory-related behavioral changes on gene-silencing of *Arg1* and *Odc1* in APP/PS1 mice, we performed Y-maze test and passive avoidance test (PAT) 3 weeks after virus injection, as described in Fig 6H. We found that low level of percentage alteration (% alteration) of APP/PS1 mice was restored by genetic silencing of astrocytic *Arg1* and *Odc1* (Fig. 6I), indicating rescue of the impaired spatial short-term memory. Furthermore, we assessed PAT, which accounts for DG-related long-term memory. One day post-acquisition, significantly short latency to dark room was observed in APP/PS1 mice, indicative of impaired long-term memory (Fig. 6J). Silencing astrocytic *Arg1* and *Odc1* was able to recover this entry latency partially, yet significantly (Fig. 6J), thereby ameliorating the long-term memory impairment of APP/PS1 mice. Previous reports point towards the correlation of aberrant tonic GABA release and memory impairment in AD (Jo et al., 2014). To understand this correlation, we plotted tonic GABA current against the PAT entry latency of each subject. Interestingly, we observed a highly negative correlation between entry latency and tonic GABA current (Fig. 6K), indicating that the higher the tonic GABA release from astrocytes, the worse the memory. Taken together, we concluded that ARG1 and ODC1 critically regulate tonic GABA levels, synaptic firing, and memory impairment in APP/PS1 mice, suggesting their potential as novel therapeutic targets for AD.

### Aβ plaques are decreased by gene-silencing of *Odc1* in APP/PS1 mice

To investigate the effect of ARG1 and ODC1 on Aβ plaques, we examined the number of neuritic plaques in APP/PS1 mice on *Arg1* and *Odc1* gene silencing. We injected lentivirus carrying pSicoR-Arg1shRNA-GFP or pSicoR-Odc1shRNA-GFP into the DG layer of WT or APP/PS1 mice and immunostained for PyrPeg, which can selectively detect neuritic plaques (Choi et al., 2020) (Fig. 7A). Four weeks after virus injection, we observed that the PyrPeg-positive plaques were significantly increased in APP/PS1 mice compared to WT mice (Fig. 7B). Interestingly, the number of PyrPeg-positive plaques was significantly reduced in *Odc1* gene-silenced group, but no significant change was observed in the *Arg1*-silenced group (Fig. 7B). Further, we employed the use of RNASeq to examine the expression levels of genes encoding for APP-processing-related proteins, such as, α-secretase, β-secretase and γ-secretase in primary astrocyte cultures treated with Aβ, with or without DFMO (Fig. 7C and D). We found that genes associated with the amyloidogenic cleavage of APP, β-secretase (*Bace1* and *Bace2*) and γ-secretase related genes (*Psen1, Psen2, Aph1b, Aph1c, Ncstn*) showed upregulated levels in Aβ-treated astrocytes compared to control (Fig. 7D). In contrast, this upregulation was reversed by DFMO treatment, accompanied by upregulation of non-amyloidogenic cleavage-related α-secretase genes (Fig. 7D). These results suggest that ODC1 inhibition promotes the non-amyloidogenic cleavage of APP over the amyloidogenic cleavage. Taken together with results from Fig. 3, these data imply that ODC1-inhibition displays a dual beneficial effect in AD-like conditions, by increasing Aβ detoxification as well as reducing the amyloidogenic load.

**Figure 7.**
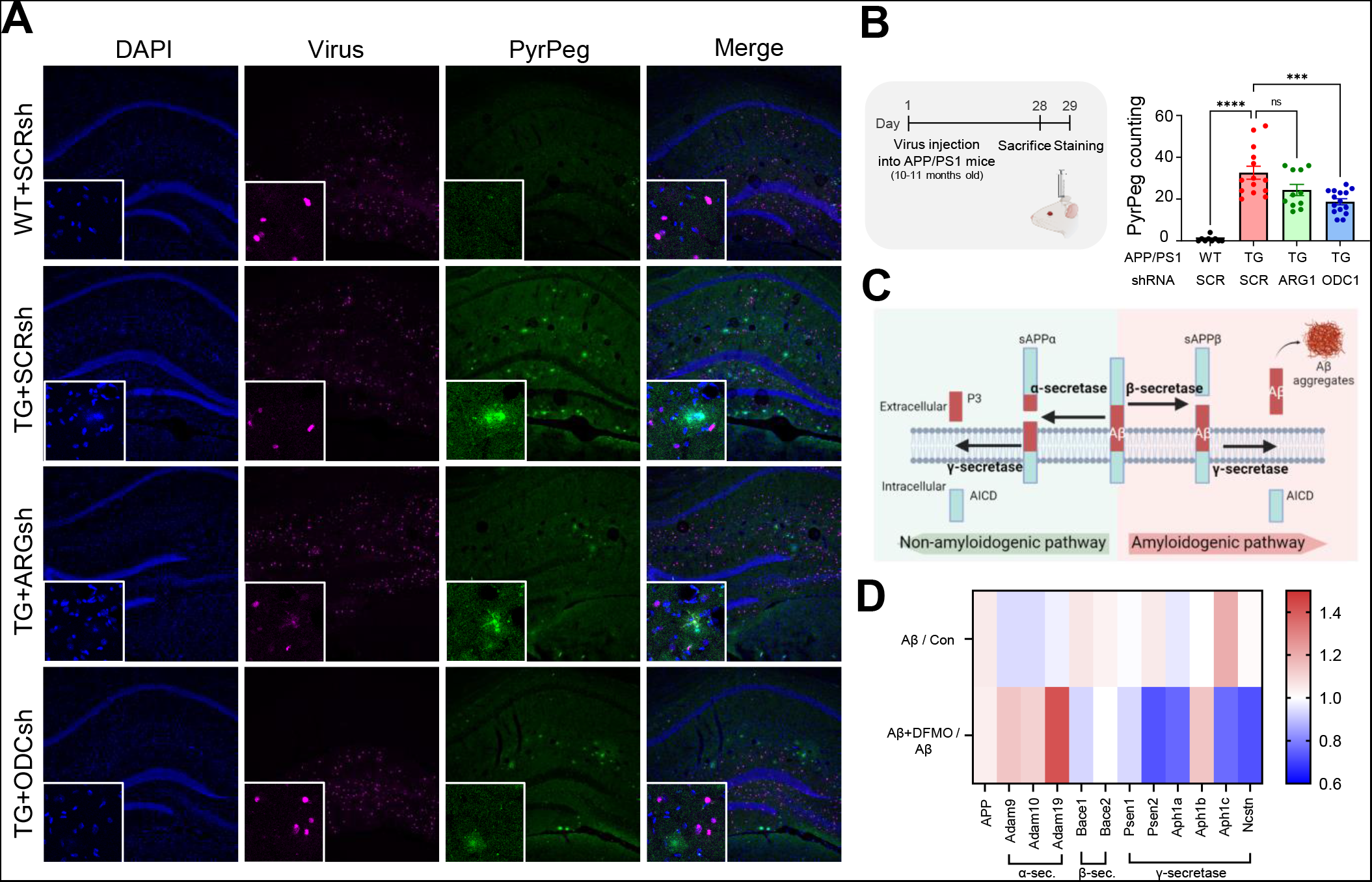
Aβ plaques are decreased by gene-silencing of ODC1 in APP/PS1 mice (see also Figure S5) **(A)** Immunostaining with DAPI and PyrPeg in Lenti-pSicoR-ScrshRNA/ARGshRNA/ODC1shRNA-GFP virus injected APP/PS1 mice (pseudo-colored). **(B)** Left, Schematic representation of experimental timeline of virus injection and IHC of APP/PS1 mice. Right, Quantification of the number of PyrPeg-positive plaques. **(C)** Schematic diagram of APP processing in amyloidogenic and non-amyloidogenic pathways. **(D)** Heat-map plot for the mRNA expression level of α-secretase, β-secretase and γ-secretase related genes. Data represents Mean ± SEM. *, p<0.05; **, p<0.01; ***, p<0.001.

## Discussion

In this study, we have delineated the detailed molecular mechanisms and metabolic pathways of the astrocytic urea cycle leading to putrescine degradation pathway in AD-like conditions. We have identified the elevated genes (*CPS1, OCT, ASL, ARG1*, and *ODC1*) and metabolites (aspartate, ammonia, urea, putrescine, GABA) in AD-associated reactive astrocytes and demonstrated that urea metabolism under normal condition is non-cycle, while it becomes fully cyclic upon Aβ-treatment. Our study unveils the comprehensive mechanisms of Aβ uptake and autophagic degradation, entrance of excess aspartate and ammonia into the urea cycle, increased putrescine and GABA production and the consequent memory impairment in AD, exacerbated by excessive toxic ammonia re-entering the urea cycle and amplifying the process. We highlight that ODC1 is the key enzyme that separates the detoxifying urea cycle and detrimental putrescine degradation pathway. Based on our findings, we propose that ODC1-inhibition turns toxic severe reactive astrocytes into Aβ-detoxifying astrocytes in AD (Fig S5).

Our study provides the first lines of evidence for the existence of the urea cycle in reactive astrocytes. Under normal conditions, however, we find that the astrocytes appear to have non-cyclic urea metabolism (Fig. 1), as seen by the minimal expression of enzymes *Cps1* and *Otc*. We found that the non-cyclic urea metabolism of healthy astrocytes switches to cyclic urea metabolism in AD-like conditions. Increased autophagic degradation of Aβ leads to accumulation of aspartate and ammonia, which can enter the urea cycle by the way of ASS1-driven arginosuccinate production and CPS1-driven carbamoyl phosphate production, respectively. Five-day treatment of Aβ was sufficient to cause aspartate accumulation (Fig. 3) and upregulate ASL1, which mediates the enzymatic breakdown of arginosuccinate to form arginine (Fig. 1). Aβ treatment was also found to accumulate ammonia (Fig. 3) and consequently turn on the expression of *Cps1* and *Otc* (Fig. 1) to facilitate entrance of toxic substrate ammonia into the urea cycle, turning on this non-cyclic-to-cyclic switch. Upregulation of urea cycle enzymes in AD-like condition without significant accumulation of intermediate metabolites (Fig. 3) implies accelerated flux of the cycle, thereby increasing build-up of end-product urea. This is consistent with previous reports of elevated urea levels in AD patients (Xu et al., 2016) and with our findings in the APP/PS1 mouse brain (Fig. 3). Our results propose that astrocytes display a unique metabolic plasticity in response to Aβ burden.

We have proposed ARG1 and ODC1 as potential therapeutic targets against AD. Between the two candidates, we propose ODC1 as a superior candidate than ARG1. ARG1 mediates the conversion of arginine to urea and ornithine, the former being the final product of the urea cycle, thereby detoxifying Aβ and ammonia. However, inhibition of ARG1 would disrupt this detoxification, ultimately limiting the clearance of toxic ammonia and Aβ, as can be seen by PyrPeg staining in Fig. 7. We find that arginine accumulates on gene-silencing of *Arg1* (Fig. 3B), which could lead to turning on of *iNOS*, production of peroxynitrites and further nitrosylation and neurodegeneration of neighboring neurons (Chun et al., 2020). Inhibition of ODC1, on the other hand, ensures maintenance of a constant cyclic flux, thereby effectively removing accumulated ammonia without the over-production of putrescine (Fig. 3C, E, G). This ensures efficient and continuous clearance of Aβ, while shutting down the detrimental putrescine degradation pathway (Fig. 7B). The accumulated urea can be efficiently removed by the blood stream, as was observed by unchanged extracellular urea concentration *in-vivo* after DFMO injection (Fig. 3K and L). Urea transporter B (UTB), a protein that mediates the removal of accumulated urea, is known to have altered expression levels in response to varying systemic urea load (Inoue et al., 2005) to facilitate the removal of accumulated urea. UTB expression has been found to increase in Huntington’s Disease patient brains in response to elevated urea (Handley et al., 2017). Consistently, our RNAseq results indicate that expression of *UTB* (*Slc14a1*) is increased in Aβ-treated astrocytes. Therefore, we conclude that ODC1 is a more effective therapeutic target compared to ARG1.

Our study not only reveals the existence of the astrocytic urea cycle, but also directs attention to its duality- beneficial role in dealing with accumulated ammonia but the detrimental effect of the downstream conversion of ornithine to putrescine, as seen in reactive astrocytes. We identify ODC1 as the molecular bridge between the two. This clarification allows us to distinctly separate the two pathways and selectively block the synthesis of putrescine, consequently reducing the production of toxic ammonia and H_2_O_2_ and preventing reactive astrogliosis. We also find that ODC1 is specifically turned on in the astrocytes of APP/PS1 mice (Fig. 5), suggesting ODC1 as a reactive astrocyte marker. Inhibition of ODC1 can therefore, selectively target reactive astrocytes to retain the positive detoxifying role of astrocytes without initiating the cascade of events that leads to H_2_O_2_-mediated neurodegeneration. Furthermore, we find that ODC1-inhibition reduces Aβ-load by switching APP processing from amyloidogenic to non-amyloidogenic (Fig. 7C and D). The detailed molecular mechanism of this switch is not known. However, we hypothesize the role of H_2_O_2_ and GABA in this switch, which requires future investigations. This disease-modifying trifecta – 1) switch from amyloidogenic to non-amyloidogenic APP processing for reducing Aβ load, 2) Aβ clearance by the urea cycle and 3) reduced neurodegeneration via H_2_O_2_ – opens the exciting possibility of ODC1-inhibition having a multifaceted therapeutic effect.

The dichotomous roles of reactive astrocytes in brain diseases have long been debated (Chun and Lee, 2018). Astrocytes, the homeostatic regulators of the CNS, strive to clear toxic molecules and respond to insult to reverse CNS pathology. In response to severe insult, however, reactive astrocytes are known to do more harm than good to the CNS. For decades, the benefits of the neuroinflammatory astrocytic response for toxin clearance and neuroprotection have been contradicting its neurotoxic effects such as increased oxidative and nitrosative stress, aberrant GABA and neurodegeneration (Chun et al., 2020; Chun and Lee, 2018). Our study identifies the underlying molecular pathways that attribute to this dual nature of reactive astrocytes and clarifies this dichotomy: the beneficial switching-on of the urea cycle to detoxify Aβ breakdown-products and the detrimental upregulation of ODC1 to augment the production of toxic H_2_O_2_ and ammonia as by-products of the putrescine degradation pathway. H_2_O_2_ contributes to hypertrophy and neurodegeneration while ammonia re-enters the urea cycle to continue the pernicious process. We propose that the co-existence of the two opposing pathways is the very nature that makes the reactive astrocytes dichotomous.

Our study also proposes that the inhibition of ODC1 prevents AD-associated symptoms. The substrate-analog DFMO, originally designed as an anti-cancer agent (LoGiudice, 2018), is the most widely used and effective ODC1-inhibitor. Several studies in the past have shown a protective effect of DFMO treatment on AD-like pathology and memory (Kan et al., 2015). However, it has been found that long-term inhibition of ODC1 proves to be a major challenge, requiring large doses of DFMO (up to 3g per kg animal weight) due to the fast rate of *in vivo* drug metabolism (Wang, 1991), and having dose-dependent side effects, such as ototoxicity (McWilliams et al., 2000). These reports seem to be contradicted by older reports, discussing the reversible and cumulative non-significance of these side effects (Gerner, 1999) when DFMO is administered as a cancer chemotherapeutic. A single-case human study involving a mild cognitive impairment patient administered with 500 mg of DFMO for 12 months showed no significant improvement in memory (Alber et al., 2018). These controversial reports indicate that DFMO may not be a suitable drug candidate to target ODC1 in AD. Our study implores the need to develop better ODC1-inhibition strategies, in the form of small-molecule inhibitors or anti-sense oligos that can circumvent the shortcomings of currently available inhibitors and effectively metabolize Aβ to urea without producing putrescine, GABA, H_2_O_2_ and more ammonia. The novel concepts and tools from this study will offer a deeper understanding of the mechanism of AD pathogenesis, providing new therapeutic targets and strategies to fight against AD.

## Author contributions

Y.H.J., M.B., S.J., J.E.O., S.Y., U.C., J.K., W.K., J.L., Y.M.P., and J.L. performed experiments. I-J.C., H.L., H.R., and C.J.L. supervised the analysis and manuscript

## Acknowledgements

This work was supported by the Institute for Basic Science (IBS), Center for Cognition and Sociality (IBS-R001-D2). This study was also supported by the National Research Foundation (NRF) Grants from the Korean Ministry of Education, Science and Technology (2018M3C7A1056894, NRF-2020M3E5D9079742) and KIST Grants (2E30954 and 2E30962).

## Supplemental figure titles and legends

**Figure S1 related to Figures 3 and 4.**
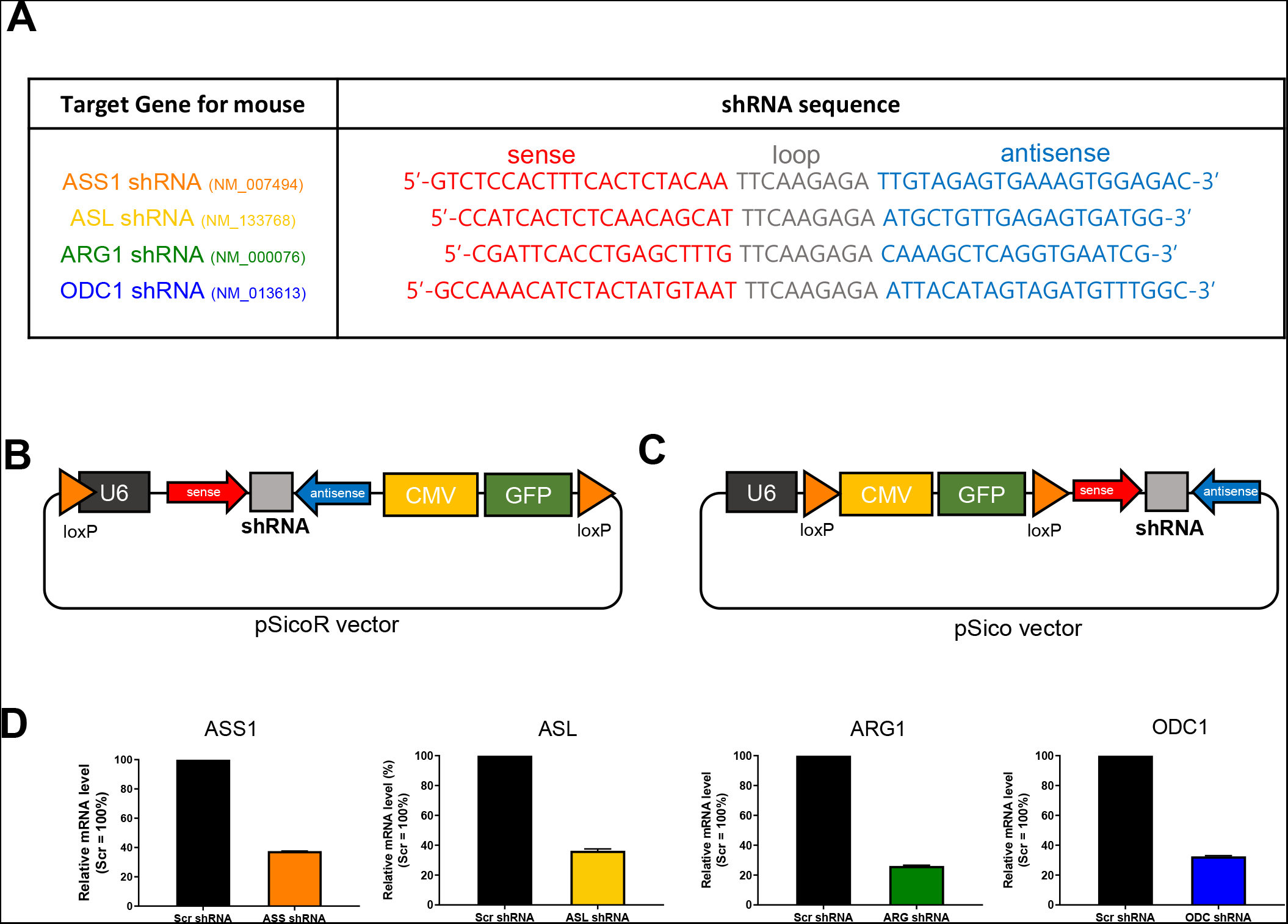
Development of shRNAs for Urea cycle enzyme. **(A)** Candidate sequences for Ass1-, Asl-, Arg1- and Odc1-shRNA. **(B, C)** pSicoR shRNA and pSico shRNA vector information. **(D)** Relative mRNA level of target gene after shRNA transfection compared to Scr-shRNA (Scr = 100%).

**Fig. S2 related to Figure 3.**
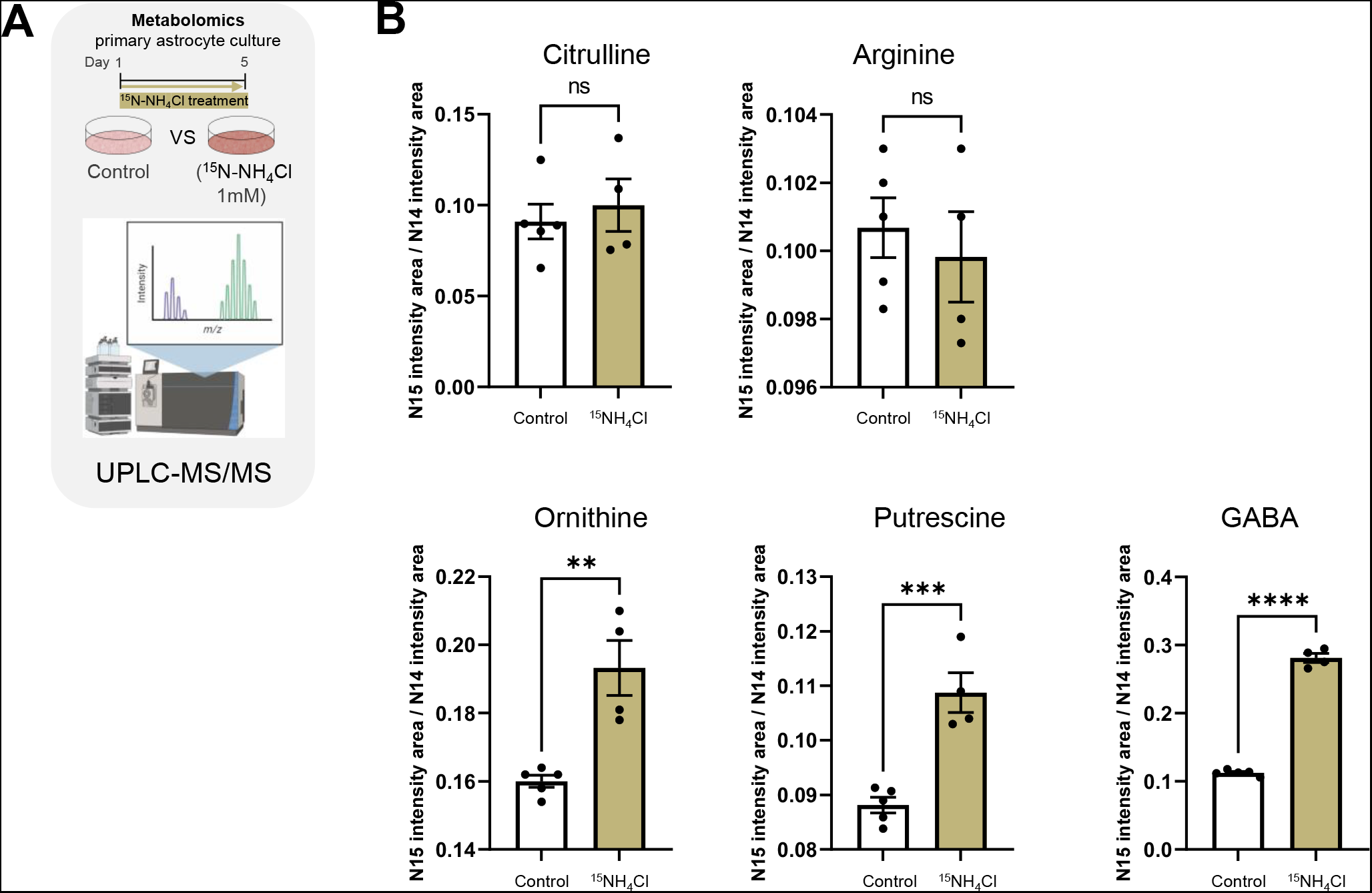
Concentration of ^15^N-containing urea cycle metabolites in ^15^NH_4_Cl-treated astrocytes. **(A**) Schematic representation of metabolomic analysis of ^15^NH_4_Cl-treated primary cultured astrocytes using UPLC-MS/MS. **(B)** Bar graphs for concentration of each metabolite from control and ^15^NH_4_Cl-treated astrocytes (Citrulline, Arginine, Ornithine, Putrescine and GABA). Individual dots represent separate batches of cell culture. Data represents Mean ± SEM. **, p < 0.01; ***, p < 0.001; ****, p<0.0001 (Student’s t-test).

**Fig. S3 related to Figure 3.**
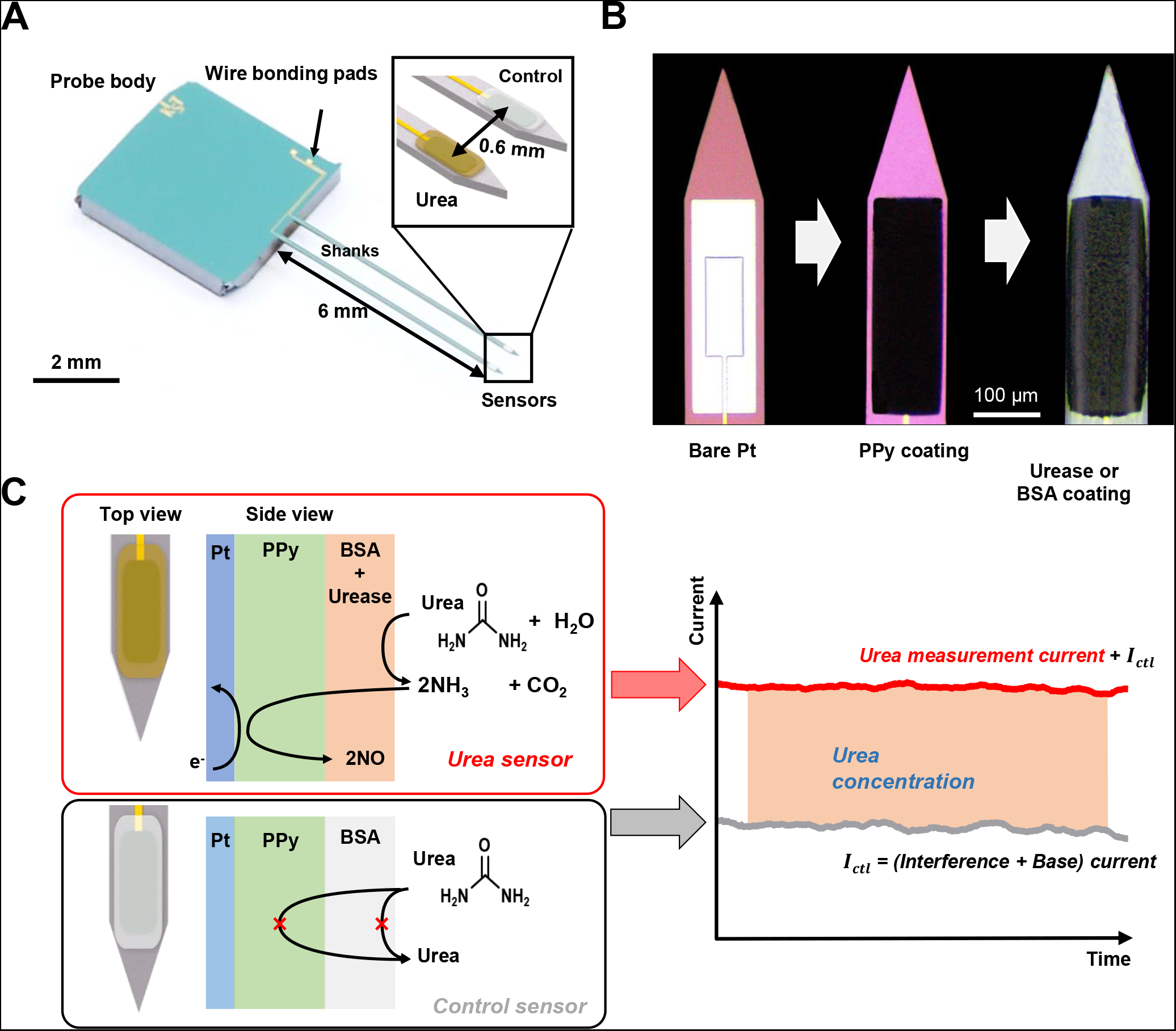
Fabrication and operating principle of the silicon probe with urea sensor to measure urea concentration *in vivo*. **(A)** Representative image of the silicon probe with urea sensor. Inset, schematic diagram of the structure. **(B)** Close-up microscope image of the biosensor electrode composed of the bare platinum (Pt) electrode, poly-pyrrole (PPy) and urease or BSA layers. **(C)** Schematic diagram illustrating the mechanism of monitoring urea concentrations using both the urea sensor and the control sensor with a urease or BSA layer on PPy. The concentration of urea is calculated by the difference between the currents from the urea sensor and the control sensor.

**Figure S4 related to Figure 4.**
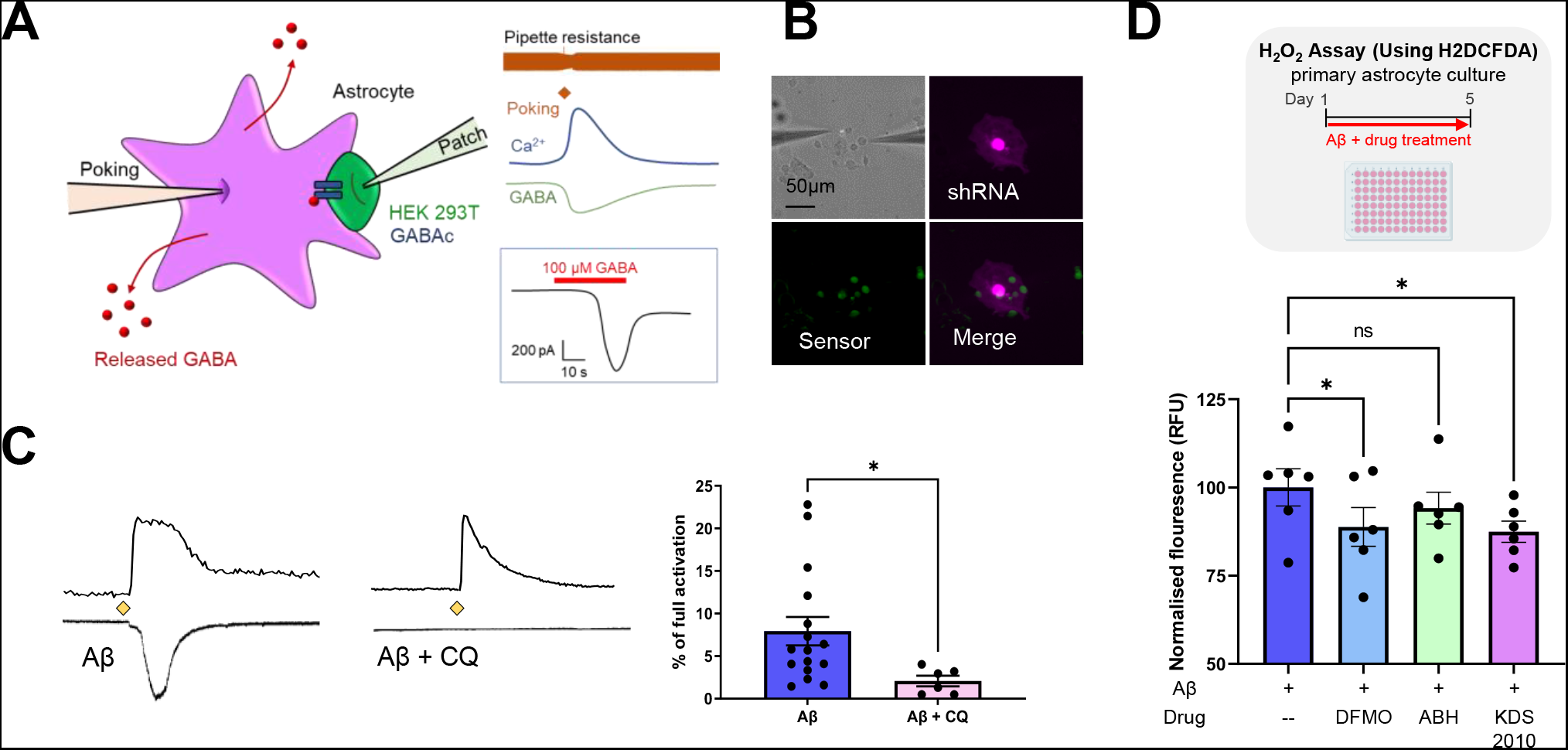
Autophagy inhibition affects GABA production. **(A)** Left, schematic diagram of Sniffer patch technique. Right, Representative traces recorded. Ca2+ signals from Fura-2 loaded astrocyte (blue trace) with poking electrode current (orange trace), whole-cell current recorded from GABAc sensor cell (green trace), and full activation of sensor cell (black trace) on application of 100μM GABA. **(B)** Representative fluorescence image for sniffer patch (scale bar, 50μm). GABA sensor cell express GFP and astrocytes transfected with shRNA express mCherry. **(C)** Representative traces and bar graph of GABA current from primary cultured astrocytes treated with Aβ with or without chloroquine (CQ). **(D)** Top, schematic representation of experimental timeline for H2DCFDA-based H_2_O_2_ detection, Bottom, Bar graph representing relative H_2_O_2_ levels in primary cultured astrocytes treated with Aβ in the presence or absence of DFMO, ABH and KDS2010. Data represents Mean ± SEM. *, p<0.01; (Student’s t-test, panel B); *, p<0.05; (2-way ANOVA, panel D).

**Figure S5 related to Figure 7.**
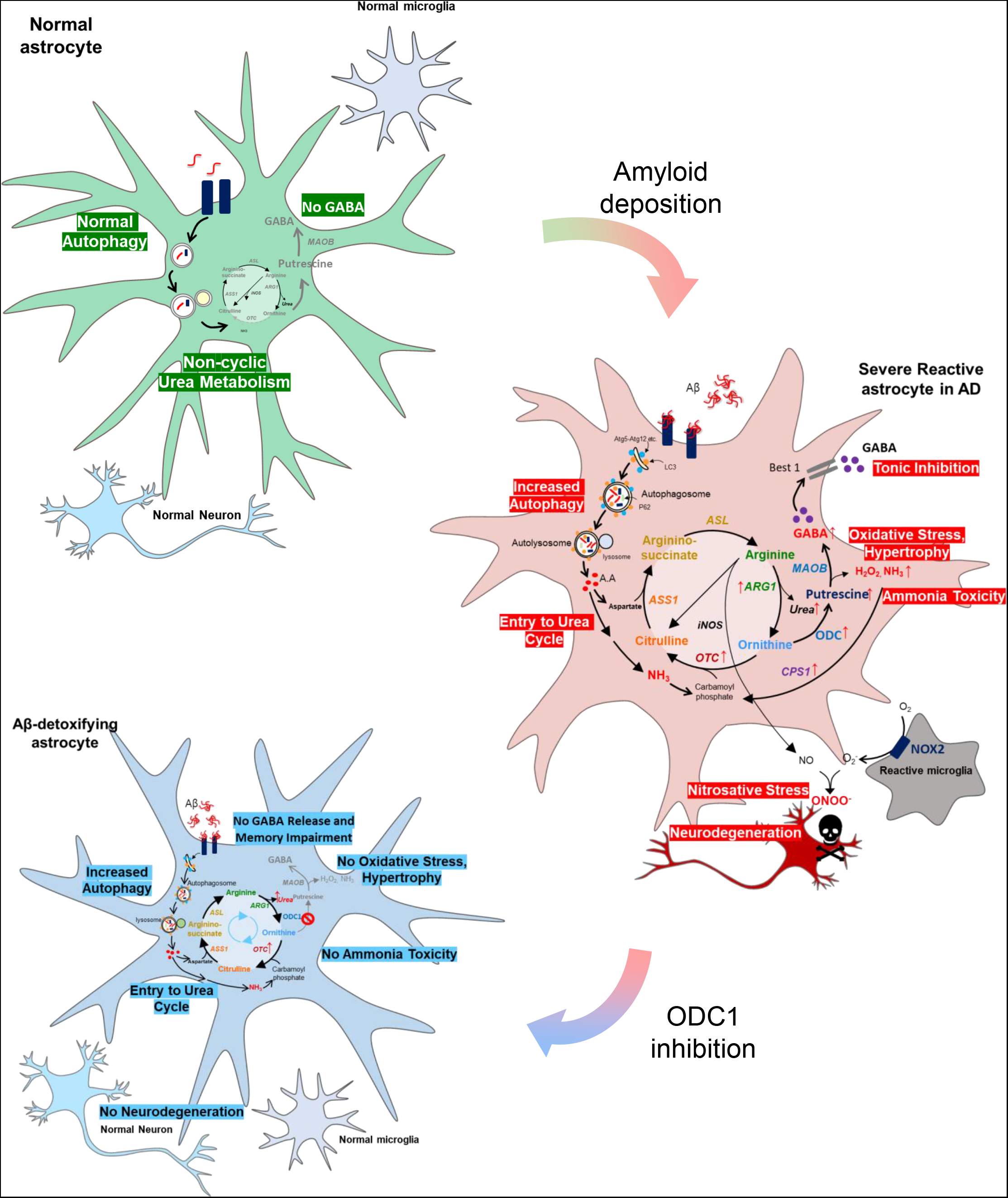
Detoxification of Aβ by the astrocytic urea cycle impairs memory in AD. Left top, in normal condition, astrocytes show normal level of autophagy and non-cyclic urea metabolism, along with low GABA level. Right middle, in AD-like condition, astrocytes show upregulation of autophagy system in response to amyloid deposition and switch-on of the urea cycle along with upregulation of enzyme ODC1, leading to increased putrescine production. MAO-B degrades putrescine to produce aberrant GABA, H_2_O_2_ and ammonia, causing tonic inhibition and oxidative stress as well as feeding ammonia back into the urea cycle, thereby continuing the detrimental process. Left bottom, ODC1 inhibition blocks the conversion of ornithine to putrescine and increases the flux of the urea cycle to produce urea. Reduced production of putrescine consequently reduces MAO-B-mediated aberrant production of GABA, H_2_O_2_ and ammonia, thereby preventing oxidative stress, neurodegeneration and memory impairment, as well as ammonia toxicity.ODC1 inhibition can, therefore, facilitate Aβ detoxification while preventing the detrimental features of astrocytic reactivity.

## STAR Methods text

### RESOURCE AVAILABILITY

#### Lead contact

Further information and requests for resources and reagents should be directed to and will be fulfilled by the lead contact Dr C Justin Lee

#### Materials availability

The sequences of the shRNAs used in this study have been provided in the Supplementary figures. The viruses used in this study were provided by and are available with the Institute for Basic Science Virus Facility (https://www.ibs.re.kr/virusfacility/, Daejeon, South Korea) upon request. The newly developed urea sensor used in this study will be available upon request.

#### Data and code availability

Next Generation RNA-seq data has been deposited at NCBI GEO and identifier will be provided when available. Microscopy data reported in this paper will be shared by the lead contact upon request. This paper does not report any original code. Any additional information required to reanalyze the data reported in this paper is available from the lead contact upon request.

### EXPERIMENTAL MODEL AND SUBJECT DETAILS

#### Animals

All experiments performed on AD mouse model were performed on APPswe/PSEN1dE9 (APP/PS1) mice of B6C3 hybrid background were used (RRID: MMRRC_034829-JAX) originated from Jackson Laboratory (USA, stock number 004462) and maintained as hemizygotes by crossing transgenic mice to B6C3 F1 mice. Genotypes were determined by PCR using the following primers – APP/PS1f - 5′ AAT AGA GAA CGG CAG GAG CA 3′; APP/PS1r - 5′ GCC ATG AGG GCA CTA ATC AT 3′. All mice were maintained under 12:12-h light-dark cycle (lights on at 8:00 AM) and had *ad libitum* access to food and water. Care and handling of animals were in accordance with the guidelines of the Institutional Animal Care and Use Committee of IBS (Daejeon, South Korea) and Korea Institute of Science and Technology (Seoul, South Korea). Slice patch, IHC and behavioral tests were performed on virus-injected mice, for which both sexes of 10- to 13-month-old transgenic mice and wild-type littermates were used.

### METHOD DETAILS

#### Cell culture

Primary cortical astrocytes were prepared from 1-day postnatal C57BL/6 mice as previously described (Woo et al., 2012). The cerebral cortex was dissected free of adherent meninges, minced, and dissociated into a single-cell suspension by trituration. Dissociated cells were plated onto plates coated with 0.1 mg/ml poly-D-lysine (Sigma). Cells were grown in Dulbecco’s modified Eagle’s medium (DMEM, Corning) supplemented with 4.5 g/L glucose, L-glutamine, sodium pyruvate, 10% heat-inactivated horse serum, 10% heat-inactivated fetal bovine serum, and 1000 units/ml of penicillin–streptomycin. Cultures were maintained at 37°C in a humidified atmosphere containing 5% CO2. Three days later, cells were vigorously washed with repeated pipetting using medium and the media was replaced to remove debris and other floating cell types.

#### Quantitative real-time RT-PCR

Quantitative real-time RT-PCR was carried out using SYBR Green PCR Master Mix as described previously (Kwak et al., 2020). Briefly, reactions were performed in duplicates in a total volume of 10μl containing 10pM primer, 4μl cDNA, and 5μl power SYBR Green PCR Master Mix (Applied Biosystems). The mRNA level of each gene was normalized to that of *Gapdh* mRNA. Fold-change was calculated using the 2-ΔΔCT method. The following sequences of primers were used for real-time RT-PCR.

GAPDH forward: 5′- ACC CAG AAG ACT GTG GAT GG -3′ GAPDH reverse: 5′-CAC ATT GGG GGT AGG AAC AC-3′.

ASL forward: 5’- GAT GCC CCA GAA GAA AAA C -3’ ASL reverse: 5’- GTG CTT GGA AGT CCC TTG AG – 3’.

ASS1 forward: 5’- CAA TGA AGA GCT GGT GAG CA -3’ ASS1 reverse: 5’-TCT GAA GGC GAT GGT ACT CC – 3’.

ARG1 forward: 5’ – CGA GGA GGG GTA GAG AAA GG -3’ ARG reverse: 5’- ACA TCA ACA AAG GCC AGG TC – 3’

ODC1 forward: 5’ – CGT CAC TCC CTT TTA CGC AG -3’ ODC1 reverse: 5’-AGA TAA CCC TCT CTG CAG GC – 3’

OTC forward: 5’ – CCA AAG GGT TAT GAG CCA GA- 3’ OTC reverse: 5’- GCT GCT TCC AGT GGA TCA TT -3’

CPS1 forward: 5’- CTG GCT GGC TAC CAA GAG TC -3’ CPS1 reverse: 5’- ATT GGG ATC CAC AAA ATC CA – 3’

AGMAT forward: 5’ – GGG TGT CTG TTC CAT GAT GC -3’ AGMAT reverse: 5’-GAT GCG ACA GGG TCC AAA TC – 3’

ADC forward: 5’ TGC ACA AGT TTC GAG GTC AC - 3’ ADC reverse: 5’ - GTG CTT GGC AGC ATA CTT GA – 3’

#### Illumina Hiseq library preparation and sequencing

Sample libraries were prepared by Ultra RNA library prep kit (NEBNEXT, #E7530), Multiplex Oligos for Illumina (NEBNEXT, #E7335) and polyA mRNA magnetic isolation module (NEBNEXT, #E74900) according to the manufacturer’s instructions. Full details of the library preparation and sequencing protocol are provided on the website (https://international.neb.com/products/e7530-nebnext-ultra-rna-library-prep-kit-for-illumina#Product%20Information). The Agilent 2100 Bioanalyzer (Agilent Technologies) and the associated High Sensitivity DNA kit (Agilent Technologies) were used to determine quality and concentration of the libraries. Sample libraries for sequencing were prepared by the HiSeq Reagent Kit Preparation Guide (Illumina, San Diego, CA, USA) as described previously (Caporaso et al 2012). Briefly, the combined sample library was diluted to 2nM, denatured with 0.2 N fresh NaOH, diluted to 20pM by addition of Illumina HT1 buffer. The library (600μl) was loaded with read 1, read 2 and index sequencing primers on a 150-cycle (2×75 paired ends) reagent cartridge (Hiseq reagent kit, Illumina), and run on a HiSeq NEXT generation high-throughput sequencer (Illumina). After the 2 × 75 bp Illumina HiSeq paired-end sequencing run, the data were base called and reads with the same barcode were collected and assigned to a sample on the instrument, which generated Illumina FASTQ files.

#### NGS data analysis

BCL files obtained from Illumina HiSeq2500 were converted to fastq and demultiplexed based on the index primer sequences. The data was imported to Partek Genomics Suite (Flow ver. 10.0.21.0328; Copyright 2009, Partek Inc., St. Louis, MO, USA), where further processing was carried out. Read quality was checked for each sample using FastQC. High-quality reads were aligned using STAR (2.7.3a). Aligned reads were quantified to the mouse genome assembly (mm10, Ensemble transcripts release 99) and normalised to the median ratio (for analysis using DeSeq2). Differential analysis was carried out using DeSeq2, pathway analysis was carried out with reference to the KEGG database.

#### Human brain samples

Normal subject and AD human brain samples were neuropathologically examined and prepared according to procedures previously established by the Boston University Alzheimer’s Disease Research Center (BUADRC). Next-of-kin provided informed consent for participation and brain donation. Institutional review board approval for ethical permission was obtained through the BUADRC center. This study was approved for exemption by the Institutional Review Board of the Boston University School of Medicine because it only included tissues collected from post-mortem subjects not classified as human subjects. The study was performed in accordance with institutional regulatory guidelines and principles of human subject protection in the Declaration of Helsinki. The sample information is listed in Supplementary Table S1.

#### Double staining immunohistochemistry for the human postmortem brain

##### First staining

Paraffin-embedded tissues were sectioned in a coronal plane at 20μm. Endogenous alkaline phosphatase was blocked using 3% hydrogen peroxide in TBS. Sections were blocked with 2.5% normal horse serum (Vector Laboratories) for 1 hr and then incubated with GFAP antibody (1:200 dilution) (Cat#AB5541, Millipore, Burlington, MA, USA) for 24 hr. After washing three times with PBS, the slides were processed with Vector ABC Kit (Vector Laboratories, Inc., Burlingame, CA, USA). The GFAP immunoreactive signals were developed with DAB chromogen (Thermo Fisher Scientific, Meridian, Rockford, IL, USA).

##### Second staining

To verify localization of ARG1, ODC1, or OTC in reactive astrocytes, ARG1, ODC1, or OTC antibody was incubated over the GFAP-stained tissue slides for 24 hrs. After washing with PBS three times, sections were incubated with ImmPRESS-AP anti-rabbit IgG (alkaline phosphatase) polymer detection reagent (Vector Laboratories: MP-5402) for 30 minutes at room temperature. ARG1, ODC1, or OTC signals were developed with a Vector Red alkaline phosphatase substrate kit (Vector Laboratories). Double- stained tissue slides were processed back to xylene through an increasing ethanol gradient [70%, 80% and 95% (1 time), and 100% (2 times)] and then mounted.

#### Metabolite analysis

For metabolite analysis, aspartate, ornithine, ^15^N-ornithine, arginine, ^15^N-arginine, citrulline, ^15^N-citrulline, glutamate, putrescine, ^15^N-putrescine, GABA and ^15^N-GABA were analyzed. The system used for the analyses was an Exion LC AD UPLC coupled with an MS/MS (Triple Quad 4500 System, AB Sciex LLC, Framingham, USA) using an Acquity® UPLC BEH C18 column (1.7 μm, 2.1 mm x 75 mm, Waters, USA) at 50°C, controlled by Analyst 1.6.2 software (AB Sciex LP, Ontario, Canada). 70% methanol (100μL, with internal standard d5-glutamine at a final concentration of 1μM) was added to the astrocyte sample pellets and vortexed for 30s. Cells were lysed by three consecutive freeze/thaw cycles using liquid nitrogen, and the lysate was centrifuged for 10 min at 20,817 x g (14,000 rpm) at 4°C. The supernatant (5 μL) from each sample was used for DNA normalization. DNA concentrations were analyzed using a Nano-MD UV-Vis spectrophotometer (Scinco, Seoul, Korea). 40μL of the supernatant from each sample was evaporated to dryness at 37°C under a gentle stream of nitrogen. Phenylisothiocyanate (PITC) derivatization was performed by adding 50μL of a mixture of 19:19:19:3 ethanol:water:pyridine:PITC (v/v) and the mixture was vortexed for 30s and shaken for 20 min. Then the mixture was evaporated to dryness at 37°C under a gentle stream of nitrogen. The residue was reconstituted by adding 50μL of the mobile phase A (0.2% formic acid in deionized water): B (0.2% formic acid in acetonitrile) = 5:5 solvent and vortexing for 30s. The initial chromatographic conditions were 100% solvent A at a flow rate of 0.4 mL.min^−1^. After 0.9min at 15% B, solvent B was set to 15% over the next 4.1min, solvent B was set to 70% over the next 5min, solvent B was set to 100% over the next 0.5min, and these conditions were retained for an additional 2min. The system was then returned to the initial conditions over the next 0.5min. The system was re-equilibrated for the next 2.5min in the initial conditions. The total running time was 15min. All samples were maintained at 4°C during the analysis, and the injection volume was 5μL. The MS analysis was performed using ESI in positive mode. The ion spray voltage and vaporizer temperature were 5.5 kV and 500°C, respectively. The curtain gas was kept at 45 psi, and the collision gas was maintained at 9 psi. The nebulizer gas was 60 psi, while the turbo gas flow rate was 70 psi. The metabolites were detected selectively using their unique multiple reaction monitoring (MRM) pairs. The following MRM mode (Q1 / Q3) was selected: arginine (m/z 310.000 / 217.000), ^15^N-arginine(m/z 311.000 / 218.000), ornithine (m/z 403.200 / 310.200), ^15^N-ornithine(m/z 404.000 / 311.200), citrulline (m/z 311.200 / 113.100), ^15^N-citrulline(m/z 313.200 / 114.100), glutamate (m/z 283.200 / 130.200), aspartate (m/z 269.200 / 116.200), putrescine (m/z 359.200 / 266.100), ^15^N-putrescine(m/z 360.200 / 267.100), GABA (m/z 238.875/ 87.103), ^15^N-GABA(m/z 239.875 / 87.103). As to monitor specific parent-to-product transitions, the standard calibration curve for each metabolite was used for absolute quantification.

#### Ammonia assay

Ammonia levels in primary astrocytes cultures were determined using an Ammonia Assay Kit (Sigma-Aldrich, #AA0100) according to kit instructions. Cell pellets were homogenized in 100μL triple-distilled water and centrifuged to clear the cell debris. The extract (in the supernatant) was assayed for ammonia concentration by measuring absorbance at 340nm using SpectraMax iD5 Multi-Mode Microplate Reader (Molecular Devices).

#### Urea assay

Urea levels in primary astrocyte cultures and APP/PS1 brain sections were determined using a Urea Assay Kit (Abcam, #83362) according to kit instructions. Approximately 20 mg of tissue was homogenized in 100μL of urea assay buffer, this extract was assayed for urea concentration. Urea was measured by OD at 570 nm using SpectraMax iD5 Multi-Mode Microplate Reader (Molecular Devices).

#### Preparation of urea biosensor probe

The silicon probe for sensing the concentration of urea is composed of two 6-mm-long shanks, and a urea sensor and a control sensor are integrated on each shank, separately. The sensing area was designed to be 100 × 300 μm^2^ and the distance between two sensors was 0.6 mm. The silicon probe with urea sensor was fabricated by the microfabrication process reported previously (Chae et al., 2021). Prior to polymerization of Polypyrole (PPy) on biosensor electrodes, the surface of the electrodes was electrochemically cleaned by applying voltage of −0.6 to 1 V with a scan rate of 100 mV/s between the electrode and a chloride-treated silver wire (Ag/AgCl) in 1x PBS for 10 times (PalmSens3, PalmSens, Houten, Provincie Utrecht, Netherlands). A PPy layer for NH3 detection was coated on both urea and control sensors. Polymerization of PPy was carried out with 200mM pyrrole in 1x PBS at pH 7.4 by holding the current density of 60 μA/mm^2^ for 300 s.

Finally, urea-detecting urease layer containing bovine serum albumin (BSA, 10 mg/ml), urease (100 U/ml), and glutaraldehyde (GA, 0.75%) was coated on the urea sensor, and a BSA layer containing BSA (10 mg/ml) and GA (0.75%) was coated on the control sensor by dropping the solutions on respective probe tips, followed by cross-linking at room temperature for 1 hour.

The principle of the urea biosensor with chronoamperometry is based on the measurement of the current generated by the reaction between PPy and NH3, the latter of which is produced through the reaction between urease and urea. During the measurement, constant potential between the PPy layer and the Ag/AgCl reference electrode is applied (Pundir et al., 2019; Scott et al., 2008). Current generated by other ions donating an electron to the electrode was measured by the control sensor and subtracted from the current measured by the urease-coated sensor. Change in urea concentration as a function of current change could, therefore, be calculated as the difference between the current measured by the urea sensor and the control sensor.

#### Measurement of urea concentrations in vivo

Prior to *in vivo* experiments, the urea biosensors were characterized in aCSF solution containing 126 mM NaCl, 24 mM NaHCO_3_, 1 mM NaH_2_PO_4_, 2.5 mM KCl, 2.5 mM CaCl_2_, 2 mM MgCl_2_, and 10 mM d-(+)-glucose, and pH 7.4. The urea concentration was gradually increased in the solution and currents were simultaneously measured at a constant potential of 0.35 V vs. Ag/AgCl at both the urea and control sensors using multiple potentiostats (STAT4000, Metrohm DropSens, Spain). Immediately after characterization and calibration, animals were anaesthetized by intraperitoneal injection of 5% urethane (400 mg/kg) and fixed in a stereotaxic frame. Part of the scalp and the skull of mice were removed, and probe shanks and reference electrodes were inserted into the brain. After the probe shank was placed in the hippocampus region of the mice (coordinates: AP −1.70, ML −0.60 and −1.20, DV −2.20; millimeter, relative to bregma), 0.35 V was applied to each sensor vs. Ag/AgCl. The biosensor was allowed to stabilized for 6000 s and the *in vivo* urea concentration was calculated by averaging the difference in current measured for 1000 s. ABH (10 mg/kg mouse body weight) or DFMO (600 mg/kg mouse body weight) was administered by subcutaneous injection and allowed to perfuse/diffuse/incubate for 7000 s while measuring real-time urea-mediated current through the biosensor. After 7000 s, urea concentration was measured by averaging the current value over 1000 s.

#### Immunostaining for confocal microscopy

Cultured primary astrocytes were fixed in PFA for 15 mins and incubated for 1 h in a blocking solution (0.3% Triton-X, 2% normal serum in 0.1 M PBS) and then immunostained with a mixture of primary antibodies in a blocking solution at 4 °C on a shaker overnight. After washing in PBS 3 times, samples were incubated with corresponding fluorescent secondary antibodies for 2 h and then washed with PBS 3 times. If needed, DAPI staining was done by the addition of DAPI solution (Pierce, 1:1,000) during the second washing step. Finally, samples were mounted with fluorescent mounting medium (Dako) and dried. A series of fluorescent images were obtained with a A1 Nikon confocal microscope and processed for further analysis using NIS-Elements (Nikon) software and ImageJ (NIH) program. Any alterations in brightness or contrast were equally applied to the entire image set. Specificity of primary antibody and immunoreaction was confirmed by omitting primary antibodies or changing fluorescent probes of the secondary antibodies.

#### Sniffer patch from primary cultured astrocytes

Primary hippocampal astrocytes were prepared from P0–P2 of C57BL/6 mice as described. On the day of sniffer patch, HEK 293T cells expressing GABAC sensor were dissociated, triturated, added onto the cover glass with cultured astrocytes, and then allowed to settle for at least 1 h before sniffer patching. After HEK cells settled, cultured astrocytes were incubated with 5 μM Fura-2AM (mixed with 1ml of external solution containing 5μl of 20% pluronic acid, Invitrogen) for 40 min and washed at room temperature and subsequently transferred to a microscope stage. External solution contained (in mM): 150 NaCl, 10 HEPES, 3 KCl, 2 CaCl_2_, 2 MgCl_2_, 5.5 glucose, pH adjusted to pH 7.3 and osmolality to 325 mOsmol kg^−1^. Intensity images of 510-nm wavelength were taken at 340-nm and 380-nm excitation wavelengths using iXon EMCCD (DV887 DCS-BV, ANDOR technology). The two resulting images were used for ratio calculations in Axon Imaging Workbench version 11.3 (Axon Instruments). During sniffer patch, astrocytic TRPA1 receptor was activated by pressure poking through a glass pipette. GABAC-mediated currents were recorded from HEK 293T cells under voltage clamp (V_h_ = −60 mV) using Axopatch 200A amplifier (Axon Instruments), acquired with pClamp 11.3. Recording electrodes (4–7 MΩ) were filled with (mM): 110 Cs-gluconate, 30 CsCl, 0.5 CaCl_2_, 10 HEPES, 4 Mg-ATP, 0.3 Na_3_-GTP and 10 BAPTA (pH adjusted to 7.3 with CsOH and osmolality adjusted to 290–310 mOsm kg^−1^ with sucrose). For simultaneous recordings, Imaging Workbench was synchronized with pClamp 11.3. To normalize differences in GABAC receptor-expression on the HEK 293T cells, 100μM of GABA in the bath was applied to maximally activate the GABAC receptors after current recording. Normalization was then accomplished by dividing the current induced by GABA released from astrocytes by the current induced by bath application of GABA. Aβ_42_ peptide (H2N-DAEFRHDSGYEVHHQKLVFFAEDVGSNKGAIIGLMVGGVVIA-COOH) was prepared as previously described (Kim et al., 2004).

#### H2O2 assay

Intracellular ROS levels were detected using cell-permeable non-florescent probe 2′,7′-Dichlorofluorescin diacetate (DCFDA; Sigma, #D6883). DCFDA is de-esterified into its fluorescent form after action of intracellular esterases and oxidation by reactive oxygen species within the cell (Ishii et al., 2017). Primary astrocyte culture was seeded onto 96-well plates (Corning) and treated with Aβ (1μM) in the presence or absence of DFMO (50μM), ABH (10μM) or KDS2010 (100nM) for 5 days. On the 5th day, cells were washed twice with Hanks Buffered Salt Solution (HBSS; Welgene, #LB-003-002) and incubated with 30μM DCFDA in HBSS at room temperature for 30 minutes in the dark. The DCFDA was replaced with HBSS and fluorescence was measured using SpectraMax iD5 Multi-Mode Microplate Reader (Molecular Devices; excitation 485nm emission 538nm).

#### Immunohistochemistry of mouse tissue sections

Mice were anaesthetized with isoflurane and perfused with 0.9% saline followed by ice-cold 4% paraformaldehyde (PFA). Excised brains were post-fixed overnight at 4°C in 4% PFA and dehydrolysed in 30% sucrose for 48hours. Coronal hippocampal sections of 30μm thickness were prepared in a cryostat and stored in storage solution at 4°C. Before staining, sections were washed in PBS and incubated for 1 hour in blocking solution (0.3% Triton X-100, 2% Donkey Serum in 0.1M PBS). Primary antibodies were added to blocking solution at desired dilution and sliced were incubated in a shaker at 4°C overnight. Unbound antibodies were washed off using PBS (3 times), followed by corresponding secondary antibody incubation (in blocking solution) for 1 hour at room temperature. Unbound antibodies were washed with PBS (3 times) and DAPI was added to PBS (1:1500 dilution) in the second step to visualize the nuclei of the cells. Sections were mounted with fluorescent mounting medium (Dako) and dried. Series of fluorescent images were obtained using Nikon A1R confocal microscope and Zeiss LSM900 microscope.

26-μm Z stack images in 2μm steps were processed for Sholl analysis using the ZEN Digital Imaging for Light Microscopy blue system (Zeiss, ver. 3.2) and ImageJ (NIH, ver. 1.52s.) software. Primary antibodies were diluted to the following amounts: anti-GFAP 1:500; anti-GABA 1:500; anti-putrescine 1:200; anti-ODC1 1:200; and PyrPeg (0.1uM). Secondary antibodies were diluted 1:500 in the blocking solution. Antibody details can be found in the Key Resources Table.

#### Sholl Analysis

Sholl analysis was performed on serially stacked and maximally projected confocal images as previously described (Chun et al., 2020). Images of brain sections immunostained with GFAP antibody were used for Sholl analysis. The Sholl analysis plugin applied in ImageJ (NIH) constructs serially concentric circles at 10μm intervals from the center of GFAP signal (soma) to the end of the most distal process of each astrocyte. The number of intercepts of GFAP-positive processes at each circle and the radius of the largest circle intercepting the astrocyte are analyzed and reported.

#### Preparation of brain slices

Mice were anaesthetized with halothane and decapitated to remove the brain. The brains were sectioned in ice-cold slicing solution (sucrose, 234 mM; KCl, 2.5 mM; MgSO_4_, 104 mM; NaH_2_PO_4_, 1.25 mM; NaHCO_3_, 24 mM; CaCl_2_-2H_2_O, 0.5 mM; and glucose, 11 mM). Horizontal slices (300 mm thick) were prepared with a vibrating-knife microtome VT1000s (Leica Microsystems, Germany). For stabilization, slices were incubated in room temperature for at least 1 h in a solution containing NaCl, 124 mM; KCl, 3 mM; MgSO_4_, 6.5 mM; NaH_2_PO_4_, 1.25 mM; NaHCO_3_, 26 mM; CaCl_2_-2H_2_O, 1 mM; and glucose, 10 mM, and simultaneously equilibrated with 95% O_2_/5% CO_2_ at 25°C. In some experiments, slices were incubated with blockers during stabilization for at least 2 h.

#### Tonic GABA recording

Brain slices were prepared from mice transgenic mice and WT littermates aged around 10- to 13-months old. Mice were deeply anesthetized with 2-bromo-2-chloro-1,1,1-trifluoroethane. After anesthetization, the brain was quickly excised from the skull and submerged in ice-cold cutting solution that contained 250 mM sucrose, 26 mM NaHCO_3_, 10 mM d-(+)-glucose, 4 mM MgCl_2_, 3 mM myo-inositol, 2.5 mM KCl, 2 mM sodium pyruvate, 1.25 mM NaH_2_PO_4_, 0.5 mM ascorbic acid, 0.1 mM CaCl_2_, and 1 mM kynurenic acid (pH 7.4). All the solutions were gassed with 95% O_2_ and 5% CO_2_. Hippocampal slices were obtained by vibrating microtome (Leica VT1000S), and 300μm thick coronal hippocampal slices were maintained at room temperature in a submerged chamber with extracellular artificial cerebrospinal fluid (ACSF) solution [126 mM NaCl, 24 mM NaHCO_3_, 1 mM NaH_2_PO_4_, 2.5 mM KCl, 2.5 mM CaCl_2_, 2 mM MgCl_2_, and 10 mM d-(+)-glucose (pH 7.4)]. Slices were incubated at room temperature for at least 1 hour before recording with gas bubbling. Slices were transferred to a recording chamber that was continuously perfused with ACSF solution (flow rate, 2 ml/min). The slice chamber was mounted on the stage of an upright Olympus microscope and viewed with a 60× water immersion objective (0.90 numerical aperture) with infrared differential interference contrast optics. Cellular morphology was visualized by a charge-coupled device camera and Imaging Workbench software (INDEC BioSystems). Whole-cell recordings were made from granule cell somata located in the DG. The holding potential was −70 mV. Pipettes (resistance 6-8 MΩ) were filled with internal solution [135 mM CsCl, 4 mM NaCl, 0.5 mM CaCl_2_, 10 mM Hepes, 5 mM EGTA, 2 mM Mg–adenosine triphosphate, 0.5 mM Na2–guanosine triphosphate, and 10 mM QX-314, pH adjusted to 7.2 with CsOH (osmolarity, 278 to 285 mOsm)]. Baseline current was stabilized with d-AP5 (50 μM) and 6-cyano-7-nitroquinoxaline-2,3-dione (20 μM) before measuring tonic current. Electrical signals were digitized and sampled at 50 μs intervals with Digidata 1440 A and a MultiClamp 700B amplifier (Molecular Devices) using pCLAMP10.2 software. Data were filtered at 2 kHz. The amplitude of tonic GABA currents was measured by the baseline shift after bicuculline (100μM) administration using the Clampfit program. Tonic current was measured from the baseline to bicuculline-treated current. Frequency and amplitude of spontaneous inhibitory postsynaptic currents before bicuculline administration were detected and measured by MiniAnalysis (Synaptosoft).

#### Virus injection

Mice were anesthetized with vaporized isoflurane and placed into stereotaxic frames (Kopf). The scalp was incised, and a hole was drilled into the skull above the dentate gyrus (anterior/posterior, −1.5mm; medial/ lateral, −1.2 or +1.2mm from bregma, dorsal/ventral, - 1.8mm from the brain surface). The virus was loaded into a glass needle and injected bilaterally into the dentate gyrus at a rate of 0.2 μlmin^−1^ for 10 min (total 2μl) using a syringe pump (KD Scientific). Virus was generated from Institute for Basic science virus facility (IBS virus facility). AAV-GFAP-mCh, Lenti-psico-Scr-GFP, Lenti-pSicoR-ODC/ARG1-GFP and Lenti-pSico-ODC1/ARG1-GFP virus was used in each experiment. Mice were used for patch-clamp and *in vivo* experiments 3 weeks after the virus injection.

#### Evoked spike probability

Evoked spike probability recordings were made from horizontal brain slices with a thickness of 300μm, as described above. Synaptically evoked spikes were triggered by a tungsten bipolar electrode placed in the outer half of the middle third dentate molecular layer. Stimulus intensity was set by 0.1-Hz stimulation of lateral perforant path fibers (100μs duration; 100- to 1000-mA intensity) via a constant current isolation unit. The evoked EPSPs (excitatory postsynaptic potentials) were recorded using glass pipette electrodes (6 to 8 MΩ), filled with intracellular solution containing 120 mM potassium gluconate, 10 mM KCl, 1 mM MgCl_2_, 0.5 mM EGTA, 40 mM Hepes (pH 7.2). Spiking probability was calculated as the ratio of the number of successful (spike-generating) stimulations to the total number of stimulations. Whole-cell patch-clamp recordings, using a Multiclamp7 amplifier (Molecular Devices), were performed from DG granule cells, visually identified with in infrared video microscopy and differential interference contrast optics. Data were collected with a MultiClamp 700B amplifier (Molecular Devices) using Clampex10 acquisition software (Molecular Devices) and digitized with Digidata 1322A (Molecular Devices). Raw data were low pass–filtered at 4 kHz and collected for offline analysis at a sampling rate of 10 kHz using pClamp10.2 software.

#### Y-maze test

The mouse was put in the middle of symmetrical Y maze with three identical arms (30cm long x 16cm high) and given free access to the arms during 8 min session. The test was video-recorded (Ethovision XT, Noldus) and the results including total number of arm entries and alternation behavior were analyzed later. The percentage of alternation was calculated as (total of alternation / total arm entries −2). An entry was recorded only when all of the animal’s four limbs were within the arm.

#### Passive avoidance behavioral test

Mice were placed in the two-compartment (light and dark) shuttle chambers with a constant current shock generator (MED Associates). On the first experimental day, a mouse was put in the light chamber for the acquisition trial. After 60 s of exploration, the door separating the light and dark compartments was raised, allowing the mouse to freely enter the dark chamber. When the mouse entered a dark chamber with all four paws, the door immediately closed, and an electric foot shock (0.5 mA, 2-s duration) was delivered through the floor grid. The mouse was then returned to the home cage and the retention trial was carried out 24 h after the acquisition trial. On the second experimental day, the mouse was placed into the light chamber again. After 60 s of exploration, the door was raised to allow the mouse to enter the dark chamber. The step-through latencies of entering the dark chamber before and after the electric shock were measured to a maximum of 300 s.

## QUANTIFICATION AND STATISTICAL ANALYSIS

All analyses were done blindly. The number of experimental samples, mean and SEM values are listed in Table S2. Numbers and individual dots refer to individual samples (individual cells, separate batches of cultured cells or animals) unless clarified otherwise in figure legends. N represents number of animals used for the experiment, while *n* refers to number of cells or culture batches. Data representation and statistical analysis was performed using GraphPad Prism (Graphpad Software). For electrophysiology, Minianalysis (Synaptosoft) and Clampfit (Molecular Devices) were used. For behavioral analysis, Ethovision XT (Noldus) was used. For image analysis, ImageJ (NIH) was used. Statistical significance was set at *p<0.5, **p>0.01, ***p<0.001 and ****p<0.0001.

### Supplemental item titles

**Table S1 related to Fig. 2.** Information of postmortem brain tissues from normal subjects and AD patients.

**Table S2.** Detailed information for statistical analysis

