## Supplementary information for "Astrocytic urea cycle detoxifies Aβ-derived ammonia while impairing memory in Alzheimer’s disease"

**Figure S1 related to Figures 3 and 4.** Development of shRNAs for Urea cycle enzyme

**Figure S2 related to Figure 3.** Concentration of $^{15}N$-containing urea cycle metabolites in $^{15}NH_4Cl$-treated astrocytes

**Figure S3 related to Figure 3.** Fabrication and operating principle of the silicon probe with urea sensor to measure urea concentration *in vivo*.

**Figure S4 related to Figure 4.** Autophagy inhibition affects GABA production.

**Figure S5 related to Figure 7.** Detoxification of Aβ by the astrocytic urea cycle impairs memory in AD

**Table S1 related to Fig. 2**. Information of postmortem brain tissues from normal subjects and AD patients.

**Table S2.** Detailed information for statistical analysis

**Supplementary Figures**

**Fig. S1 related to Figures 3 and 4**

**A**

| Target Gene for mouse | shRNA sequence (sense / loop / antisense) |
|---|---|
| ASS1 shRNA (NM_007494) | 5′-GTCTCCACTTTCACTCTACAA TTCAAGAGA TTGTAGAGTGAAAGTGGAGAC-3′ |
| ASL shRNA (NM_133768) | 5′-CCATCACTCTCAACAGCAT TTCAAGAGA ATGCTGTTGAGAGTGATGG-3′ |
| ARG1 shRNA (NM_000076) | 5′-CGATTCACCTGAGCTTTG TTCAAGAGA CAAAGCTCAGGTGAATCG-3′ |
| ODC1 shRNA (NM_013613) | 5′-GCCAAACATCTACTATGTAAT TTCAAGAGA ATTACATAGTAGATGTTTGGC-3′ |

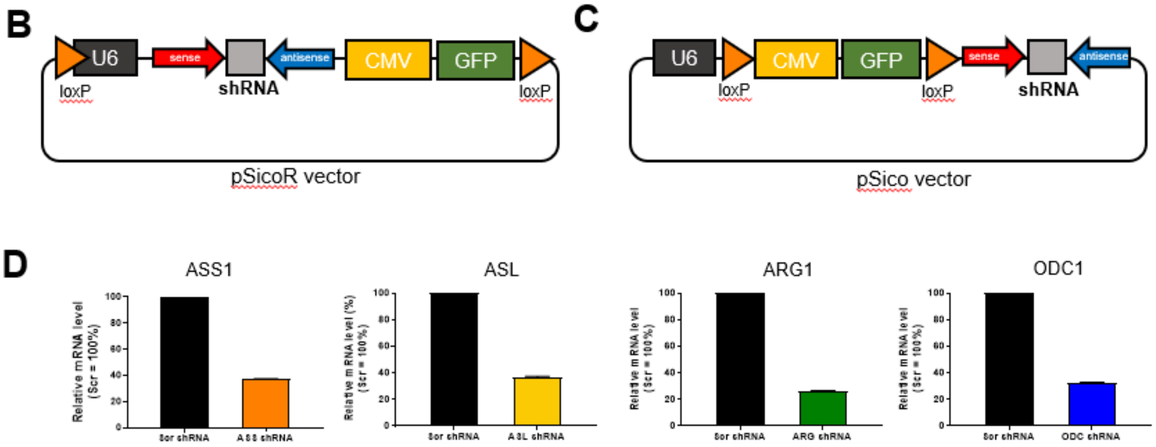


**Figure S1 related to Figures 3 and 4. Development of shRNAs for Urea cycle enzyme**

**(A)** Candidate sequences for Ass1-, Asl-, Arg1- and Odc1-shRNA. **(B, C)** pSicoR shRNA and pSico shRNA vector information. **(D)** Relative mRNA level of target gene after shRNA transfection compared to Scr-shRNA (Scr = 100%).

**Fig. S2 related to Figure 3**

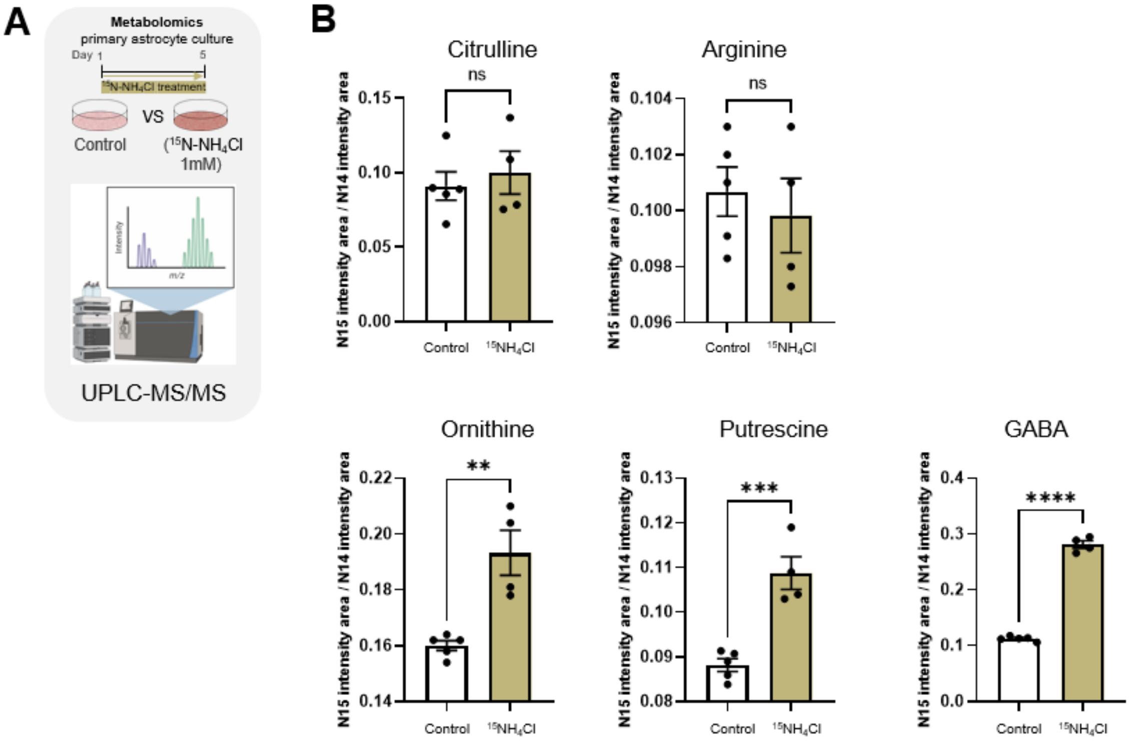


**Figure S2 related to Figure 3. Concentration of $^{15}$N-containing urea cycle metabolites in $^{15}NH_4Cl$-treated astrocytes**

**(A)** Schematic representation of metabolomic analysis of $^{15}NH_4Cl$-treated primary cultured astrocytes using UPLC-MS/MS. **(B)** Bar graphs for concentration of each metabolite from control and $^{15}NH_4Cl$-treated astrocytes (Citrulline, Arginine, Ornithine, Putrescine and GABA). Individual dots represent separate batches of cell culture. Data represents Mean ± SEM. **, $p < 0.01$; ***, $p < 0.001$; ****, $p<0.0001$ (Student's t-test).

### Fig. S3 related to Figure 3

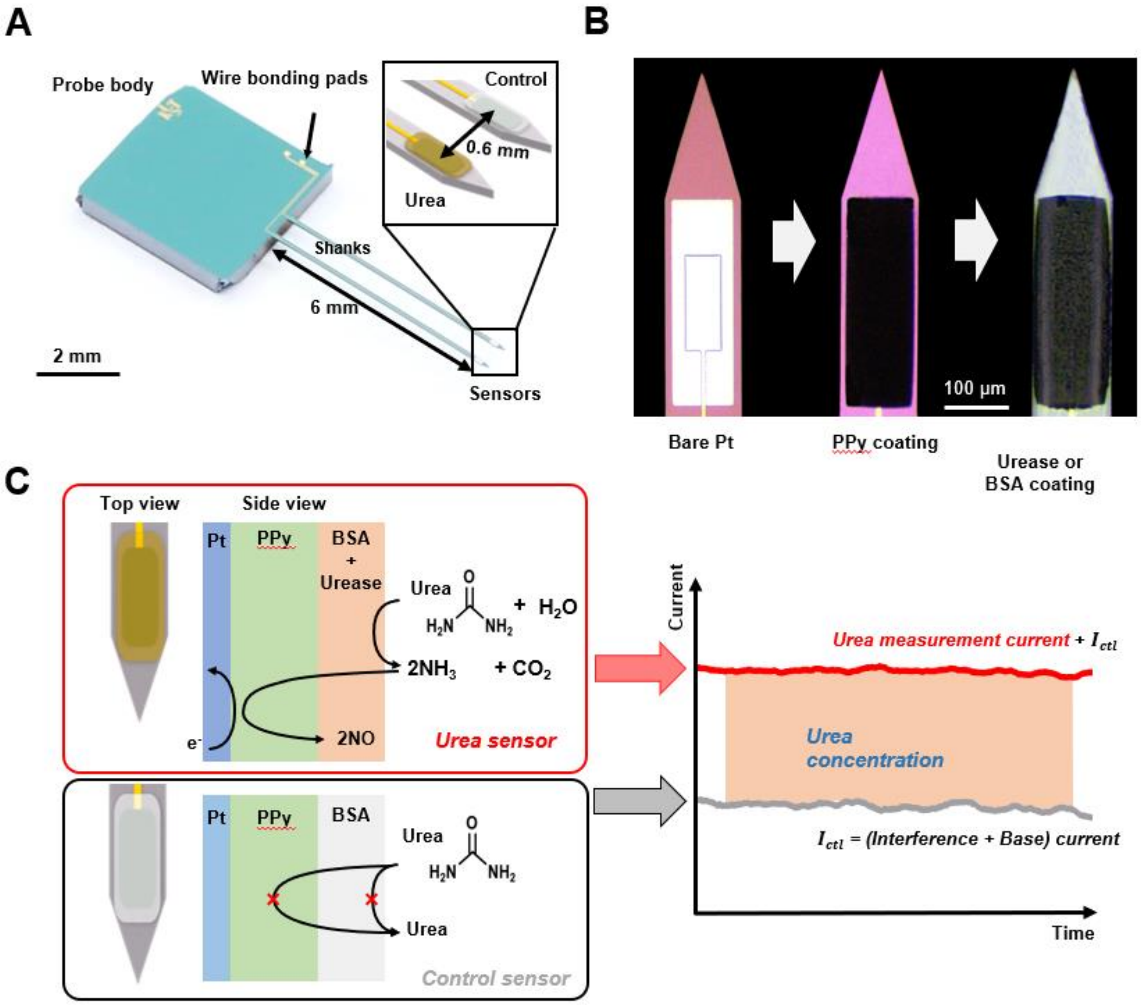


**Figure S3 related to Figure 3. Fabrication and operating principle of the silicon probe with urea sensor to measure urea concentration *in vivo*.**

**(A)** Representative image of the silicon probe with urea sensor. Inset, schematic diagram of the structure. **(B)** Close-up microscope image of the biosensor electrode composed of the bare platinum (Pt) electrode, poly-pyrrole (PPy) and urease or BSA layers. **(C)** Schematic diagram illustrating the mechanism of monitoring urea concentrations using both the urea sensor and the control sensor with a urease or BSA layer on PPy. The concentration of urea is calculated by the difference between the currents from the urea sensor and the control sensor.

Fig. S4 related to Figure 4

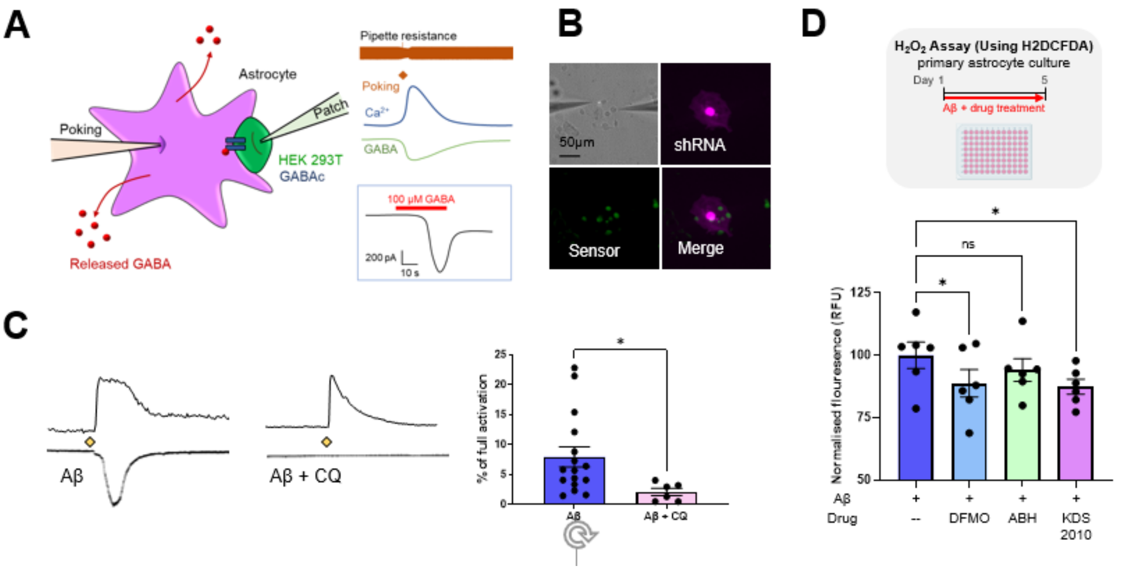


**Figure S4 related to Figure 4. Autophagy inhibition affects GABA production.**

**(A)** Left, schematic diagram of Sniffer patch technique. Right, Representative traces recorded. Ca2+ signals from Fura-2 loaded astrocyte (blue trace) with poking electrode current (orange trace), whole-cell current recorded from $GABA_c$ sensor cell (green trace), and full activation of sensor cell (black trace) on application of 100μM GABA**.** **(B)** Representative fluorescence image for sniffer patch (scale bar, 50μm). GABA sensor cell express GFP and astrocytes transfected with shRNA express mCherry**.** **(C)** Representative traces and bar graph of GABA current from primary cultured astrocytes treated with Aβ with or without chloroquine (CQ). **(D)** Top, schematic representation of experimental timeline for H2DCFDA-based $H_2O_2$ detection, Bottom, Bar graph representing relative $H_2O_2$ levels in primary cultured astrocytes treated with Aβ in the presence or absence of DFMO, ABH and KDS2010. Data represents Mean ± SEM. *, $p<0.01$; (Student's t-test, panel B); *, $p<0.05$; (2-way ANOVA, panel D)

Fig. S5

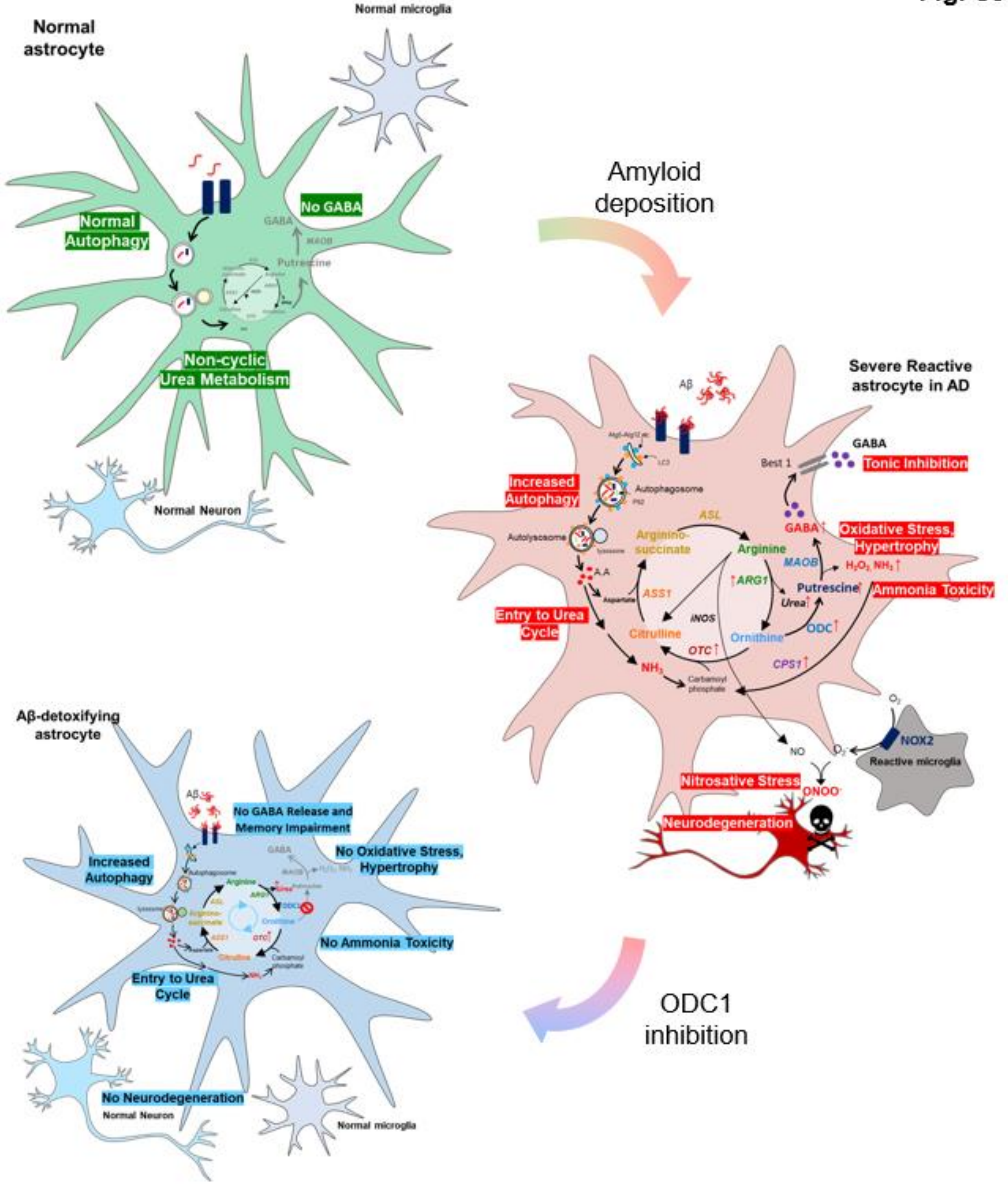


**Figure S5 related to Figure 7. Detoxification of Aβ by the astrocytic urea cycle impairs memory in AD**

Left top, in normal condition, astrocytes show normal level of autophagy and non-cyclic urea metabolism, along with low GABA level. Right middle, in AD-like condition, astrocytes show upregulation of autophagy system in response to amyloid deposition and switch-on of the urea cycle along with upregulation of enzyme ODC1, leading to increased putrescine production. MAO-B degrades putrescine to produce aberrant GABA, $H_2O_2$ and ammonia, causing tonic inhibition and oxidative stress as well as feeding ammonia back into the urea cycle, thereby continuing the detrimental process. Left bottom, ODC1 inhibition blocks the conversion of ornithine to putrescine and increases the flux of the urea cycle to produce urea. Reduced production of putrescine consequently reduces MAO-B-mediated aberrant production of GABA, $H_2O_2$ and ammonia, thereby preventing oxidative stress, neurodegeneration and memory impairment, as well as ammonia toxicity.ODC1 inhibition can, therefore, facilitate Aβ detoxification while preventing the detrimental features of astrocytic reactivity.

**Table S1 (Related to Fig. 2)**

**Information of postmortem brain tissues from normal subjects and AD patients.**

| Number | Case | Age | Sex | Braak stage |
|---|---|---|---|---|
| 1 | Normal | 87 | Female | I |
| 2 | Normal | 82 | Male | I |
| 3 | Normal | 78 | Female | I |
| 4 | Normal | 70 | Male | I |
| 5 | Normal | 82 | Male | I |
| 1 | AD | 85 | Male | V |
| 2 | AD | 87 | Female | V |
| 3 | AD | 82 | Male | V |
| 4 | AD | 79 | Female | VI |
| 5 | AD | 70 | Male | VI |

### Table S2. Detailed information for statistical analysis

| Figure No. | Result from statistical analysis |
|---|---|
| 1F | **CPS1**:<br>0h (1.000 ± 0.000, *n*=5), 6h (0.7017 ± 0.0.07649, *n*=2), 1d (1.498 ± 0.02006, *n*=2), 5d (1.488 ± 0.1797, *n*=5)<br>Unpaired Welch's t-test<br>0h vs. 1d, *p* = 0.0256; 0h vs. 5d, *p* = 0.0532. |
| | **OTC**:<br>0h (1.000 ± 0.000, *n*=6), 6h (1.408 ± 0.1878, *n*=4), 1d (1.760 ± 0.2335, *n*=5), 5d (0.8869 ± 0.1043, *n*=7)<br>Unpaired Welch's t-test<br>0h vs. 6h, *p* = 0.118; 0h vs. 1d, *p* = 0.0312; 0h vs. 5d, *p* = 0.32. |
| | **ASS1**:<br>0h (1.000 ± 0.000, *n*=5), 6h (1.977 ± 0.9395, *n*=4), 1d (1.795 ± 0.6522, *n*=6), 5d (3.294 ± 0.8366, *n*=5)<br>Unpaired Welch's t-test<br>0h vs. 6h, *p* = 0.3749; 0h vs. 1d, *p* = 0.2733; 0h vs. 5d, *p* = 0.0518. |
| | **ASL**:<br>0h (1.000 ± 0.000, *n*=5), 6h (2.000 ± 0.8498, *n*=4), 1d (2.713 ± 0.5592, *n*=7), 5d (4.628 ± 0.6781, *n*=4)<br>Unpaired Welch's t-test<br>0h vs. 6h, *p* = 0.3243; 0h vs. 1d, *p* = 0.0221; 0h vs. 5d, *p* = 0.0128. |
| | **ARG1**:<br>0h (1.000 ± 0.000, *n*=5), 6h (0.5035 ± 0.06659, *n*=4), 1d (1.554 ± 0.2419, *n*=3), 5d (1.353 ± 0.08955, *n*=8)<br>Unpaired Welch's t-test<br>0h vs. 6h, *p* = 0.0050; 0h vs. 1d, *p* = 0.1494; 0h vs. 5d, *p* = 0.0056. |
| | **ODC1**:<br>0h (1.000 ± 0.000, *n*=5), 6h (1.137 ± 0.0656, *n*=2), 1d (1.240 ± 0.0259, *n*=5), 5d (1.338 ± 0.0595, *n*=5)<br>Unpaired Welch's t-test<br>0h vs. 6h, *p* = 0.2851; 0h vs. 1d, *p* = 0.0008; 0h vs. 5d, *p* = 0.0048. |

| | |
|---|---|
| 2B | **CPS1**:<br>Normal (0.09391 ± 0.01348, *n*=8), AD (0.1930 ± 0.02652, *n*=8)<br>Unpaired two-tailed t-test, $p = 0.0050$ |
| | **OTC**:<br>Normal (0.0135 ± 0.0041, *n*=8), AD (0.0423 ± 0.0114, *n*=8)<br>Unpaired two-tailed t-test, $p = 0.0326$ |
| | **ASS1**:<br>Normal (5.109 ± 0.2703, *n*=8), AD (3.545 ± 0.3699, *n*=8)<br>Unpaired two-tailed t-test, $p = 0.0042$ |
| | **ASL**:<br>Normal (1.019 ± 0.1188, *n*=8), AD (0.5405 ± 0.05216, *n*=8)<br>Unpaired two-tailed t-test, $p = 0.0024$ |
| | **ARG1**:<br>Normal (0.0214 ± 0.0067, *n*=8), AD (0.0779 ± 0.0434, *n*=8)<br>Unpaired two-tailed t-test, $p = 0.2201$ |
| | **ODC1**:<br>Normal (2.284 ± 0.1779, *n*=8), AD (3.851 ± 0.6157, *n*=8)<br>Unpaired two-tailed t-test, $p = 0.0283$ |
| 2D | **OTC**<br>CA1: Normal [1.000 ± 0.0420, N=3; cell counting, n=30 (10 cells/case)], AD [4.078 ± 0.0719, N=3; cell counting, n=30 (10 cells/case)]<br>Unpaired two-tailed t-test, $p = 2.96324E-36$<br>CA2: Normal [1.000 ± 0.0311, N=3; cell counting, n=30 (10 cells/case)]), AD [3.174 ± 0.0783, N=3; cell counting, n=30 (10 cells/case)]<br>Unpaired two-tailed t-test, $p = 1.14232E-25$<br>CA3: Normal [1.000 ±0.0350, N=3; cell counting, n=30 (10 cells/case)]), AD [3.231 ± 0.0615, N=3; cell counting, n=30 (10 cells/case)]<br>Unpaired two-tailed t-test, $p = 8.57314E-33$<br>DG: Normal [1.000 ± 0.0539, N=3; cell counting, n=30 (10 cells/case)]), AD [2.753 ± 0.0638, N=3; cell counting, n=30 (10 cells/case)]<br>Unpaired two-tailed t-test, $p = 2.54485E-28$ |
| 2F | **ARG1**<br>CA1: Normal [1.000 ± 0.0491, N=3; cell counting, n=30 (10 cells/case)], AD (2.218 ± 0.0546, N=3; cell counting, n=30 (10 cells/case)]<br>Unpaired two-tailed t-test, $p = 1.44187E-23$<br>CA2: Normal [1.000 ± 0.0464, N=3; cell counting, n=30 (10 cells/case)]), AD (2.081 ± 0.0629, N=3; cell counting, n=30 (10 cells/case)]<br>Unpaired two-tailed t-test, $p = 2.84444E-19$<br>CA3: Normal [1.000 ±0.0370, N=3; cell counting, n=30 (10 cells/case)]), AD (2.040 ± 0.0379, N=3; cell counting, n=30 (10 cells/case)]<br>Unpaired two-tailed t-test, $p = 2.73542E-27$<br>DG: Normal [1.000 ± 0.0589, N=3; cell counting, n=30 (10 cells/case)]), AD (2.496 ± 0.0603, N=3; cell counting, n=30 (10 cells/case)]<br>Unpaired two-tailed t-test, $p = 3.99119E-25$ |
| 2H | **ODC1** |

| | |
|---|---|
| | CA1: Normal [1.000 ± 0.0423, N=3; cell counting, n=30 (10 cells/case)]), AD (2.205 ± 0.0431, N=3; cell counting, n=30 (10 cells/case)]<br>Unpaired two-tailed t-test, $p$ = 1.14923E-27<br>CA2: Normal [1.000 ± 0.0564, N=3; cell counting, n=30 (10 cells/case)]), AD (2.447 ± 0.0562, N=3; cell counting, n=30 (10 cells/case)]<br>Unpaired two-tailed t-test, $p$ = 1.27455E-25<br>CA3: Normal [1.000 ±0.0385, N=3; cell counting, n=30 (10 cells/case)]), AD (2.636 ± 0.0563, N=3; cell counting, n=30 (10 cells/case)]<br>Unpaired two-tailed t-test, $p$ = 1.65119E-29<br>DG: Normal [1.000 ± 0.0649, N=3; cell counting, n=30 (10 cells/case)]), AD (2.747 ± 0.0631, N=3; cell counting, n=30 (10 cells/case)]<br>Unpaired two-tailed t-test, $p$ = 6.4842E-27 |
| 3A | **Aspartate**:<br>Control (0.4142 ± 0.0205, $n$=5), Aβ (0.4789 ± 0.0090, $n$=5)<br>Unpaired two-tailed t-test, $p$ = 0.0204 |
| | **Putrescine**:<br>Control (11.00 ± 0.3934, $n$=6), Aβ (13.89 ± 0.4499, $n$=6)<br>Unpaired two-tailed t-test, $p$ = 0.0007 |
| | **GABA**:<br>Control (0.2850 ± 0.0135, $n$=6), Aβ (0.3957 ± 0.0191, $n$=6)<br>Unpaired two-tailed t-test, $p$ = 0.0008 |
| | **Glutamate**:<br>Control (2.951 ± 0.0786, $n$=6), Aβ (3.091 ± 0.1577, $n$=6)<br>Unpaired two-tailed t-test, $p$ = 0.4518 |
| | **Citrulline**:<br>Control (0.4000 ± 0.0315, $n$=6), Aβ (0.3948 ± 0.0242, $n$=6)<br>Unpaired two-tailed t-test, $p$ = 0.8992 |
| | **Arginine**:<br>Control (45.92 ± 5.122, $n$=6), Aβ (51.20 ± 4.348, $n$=6)<br>Unpaired two-tailed t-test, $p$ = 0.4499 |
| | **Ornithine**:<br>Control (0.2563 ± 0.0226, $n$=6), Aβ (0.2748 ± 0.0349, $n$=6)<br>Unpaired two-tailed t-test, $p$ = 0.6665 |
| 3B | **Arginine**:<br>SCR shRNA (0.8357 ± 0.1020, $n$=5), ARG1 shRNA (1.685 ± 0.1741, $n$=5), ODC1 shRNA (1.067 ± 0.1996, $n$=5)<br>One-way ANOVA with Tukey's multiple comparisons test<br>F(2, 12) = 7.170, $p$ = 0.0089<br>SCR shRNA vs. ARG1 shRNA, $p$ = 0.0084; SCR shRNA vs. ODC1 shRNA, $p$ = 0.5909 |
| | **Ornithine**:<br>SCR shRNA (0.2720 ± 0.0195, $n$=5), ARG1 shRNA (0.1876 ± 0.0148, $n$=5), ODC1 shRNA (0.4102 ± 0.0320, $n$=5)<br>One-way ANOVA with Tukey's multiple comparisons test<br>F(2, 12) = 23.23, $p$ < 0.0001 |

| | |
|---|---|
| | SCR shRNA vs. ARG1 shRNA, $p = 0.0604$; SCR shRNA vs. ODC1 shRNA, $p = 0.0033$ |
| | **Citrulline**:<br>SCR shRNA (0.0545 ± 0.0087, $n$=5), ARG1 shRNA (0.0274 ± 0.0041, $n$=5), ODC1 shRNA (0.0673 ± 0.0129, $n$=5)<br>One-way ANOVA with Tukey's multiple comparisons test<br>$F(2, 12) = 4.765$, $p = 0.0300$<br>SCR shRNA vs. ARG1 shRNA, $p = 0.1420$; SCR shRNA vs. ODC1 shRNA, $p = 0.6084$ |
| | **Glutamate**:<br>SCR shRNA (3.091 ± 0.1577, $n$=5), ARG1 shRNA (2.672 ± 0.1167, $n$=5), ODC1 shRNA (4.522 ± 0.4572, $n$=5)<br>One-way ANOVA with Tukey's multiple comparisons test<br>$F(2, 12) = 11.41$, $p = 0.0017$<br>SCR shRNA vs. ARG1 shRNA, $p = 0.5722$; SCR shRNA vs. ODC1 shRNA, $p = 0.0108$ |
| | **Aspartate**:<br>SCR shRNA (0.4789 ± 0.0090, $n$=5), ARG1 shRNA (0.6200 ± 0.0848, $n$=5), ODC1 shRNA (0.5757 ± 0.0619, $n$=5)<br>One-way ANOVA with Tukey's multiple comparisons test<br>$F(2, 12) = 1.405$, $p = 0.2829$<br>SCR shRNA vs. ARG1 shRNA, $p = 0.2678$; SCR shRNA vs. ODC1 shRNA, $p = 0.5179$ |
| 3C | **Putrescine**:<br>SCR shRNA (1.287 ± 0.0791, $n$=6), ASS1 shRNA (0.7983 ± 0.1548, $n$=6), ASL shRNA (0.8165 ± 0.0859, $n$=6), ARG1 shRNA (0.5088 ± 0.1073, $n$=6), ODC1 shRNA (0.6217 ± 0.0830, $n$=6)<br>One-way ANOVA with Tukey's multiple comparisons test<br>$F(4, 25) = 7.885$, $p = 0.0003$<br>SCR shRNA vs. ASS1 shRNA, $p = 0.0112$; SCR shRNA vs. ASL shRNA, $p = 0.0150$;<br>SCR shRNA vs. ARG1 shRNA, $p < 0.0001$; SCR shRNA vs. ODC1 shRNA, $p = 0.0006$. |
| 3E | Control (1.000 ± 0.0353, $n$=7), Aβ (1.913 ± 0.2099, $n$=9), Aβ + KDS2010 (1.243 ± 0.1802, $n$=5), Aβ + CQ (1.487 ± 0.09237, $n$=6), Aβ + ABH (1.253 ± 0.2234, $n$=8), Aβ + DFMO (1.160 ± 0.125, $n$=11)<br>Unpaired two-tailed t-test<br>Control vs. Aβ, $p<0.0001$; Aβ vs Aβ + KDS2010 $p = 0.0212$; Aβ vs. Aβ + CQ, $p = 0.0485$; Aβ vs. Aβ + ABH, $p = 0.0216$; Aβ vs Aβ + DFMO, $p = 0.0063$ |
| 3F | Control (0.1383 ± 0.0130, $n$=15), Arginine (0.3024 ± 0.0211, $n$=15), Arginine+ABH (0.2073 ± 0.0201, $n$=14)<br>One-way ANOVA with Tukey's multiple comparisons test<br>$F(2, 41) = 20.51$, $p < 0.0001$<br>Control vs. Arginine, $p < 0.0001$; Arginine vs. Arginine+ABH, $p = 0.0022$ |
| 3G | Control (0.1199 ± 0.0164, $n$=11), Aβ (0.2191 ± 0.0170, $n$=12), Aβ+DFMO (0.3871 ± 0.0316, $n$=6)<br>One-way ANOVA with Tukey's multiple comparisons test<br>$F(2, 26) = 36.78$, $p < 0.0001$<br>Control vs. Aβ, $p = 0.0018$; Aβ vs. Aβ+DFMO, $p < 0.0001$ |
| 3H | WT (1.796 ± 0.3130, n=8), APP/PS1 (2.565 ± 0.1425, n=12)<br>Unpaired two-tailed t-test, $p = 0.0223$ |
| 3L | WT (5.500 ± 0.02799, N=4), APP/PS1 (7.200 ± 0.5019, N=7), APP/PS1+ABH (3.540 ± 0.4675, N=5), APP/PS1+DFMO (6.700 ± 1.100, N=2)<br>Unpaired two-tailed t test<br>WT vs APP/PS1, $p = 0.0401$ |

| | |
|---|---|
| | Paired two-tailed t test<br>APP/PS1 vs APP/PS1+ABH, $p = 0.001$<br>APP/PS1 vs APP/PS1+DFMO, $p = 0.5$ |
| 4C | SCR(-) (3886 ± 125.8, *n*=33), SCR(+) (5924 ± 287.4, *n*=38), ASS1(+) (4448 ± 209.5, *n*=23), ASL(+) (3723 ± 75.09, *n*=30), ARG1(+) (4067 ± 143, *n*=48), ODC1(+) (3959 ± 110.3, *n*=38)<br>One-way ANOVA with Tukey's multiple comparisons test<br>$F(5, 204) = 22.48$, $p < 0.0001$<br>SCR(-) vs. SCR(+), $p < 0.0001$; SCR(+) vs. ASS1(+), $p < 0.0001$; SCR(+) vs. ASL(+), $p < 0.0001$; SCR(+) vs. ASRG1(+), $p < 0.0001$; SCR(+) vs. ODC1(+), $p < 0.0001$ |
| 4D | Naïve (0.0600 ± 0.0346, *n*=8), Aβ (9.981 ± 2.129, *n*=8)<br>Unpaired two-tailed t-test, $p = 0.0004$ |
| 4E | Scr shRNA (6.686 ± 1.210, n=9), ARG1 shRNA (0.6983 ± 0.3382, n=6), ODC1 shRNA (0.6617 ± 0.4320, n=6)<br>One-way ANOVA with Dunnett's multiple comparisons test<br>$F(2, 18) = 14.58$, $p = 0.0002$<br>Scr shRNA vs. ARG1 shRNA, $p = 0.0006$; Scr shRNA vs. ODC1 shRNA, $p = 0.0005$ |
| 4F | Control (3.785 ± 1.021, *n*=17), ABH (0.3050 ± 0.1509, *n*=12)<br>Unpaired two-tailed t-test, $p = 0.0086$ |
| 4G | Scr shRNA (7.596 ± 1.460, *n*=7), ARG1 shRNA (0.9200 ± 0.5784, *n*=7)<br>Unpaired two-tailed t-test, $p = 0.0011$ |
| 4H | Control (5.126 ± 1.569, *n*=8), DFMO (0.2040 ± 0.1005, *n*=10)<br>Unpaired two-tailed t-test, $p = 0.0028$ |
| 4I | Scr shRNA (18.84 ± 4.184, n=*9*), ODC1 shRNA (0.7014 ± 0.6183, *n*=7)<br>Unpaired two-tailed t-test, $p = 0.0021$ |
| 4J | Naïve (0.0600 ± 0.0346, *n*=8), Citrulline (3.521 ± 2.099, n=8), Aspartate (6.166 ± 1.176, *n*=10), Aspartate + Citrulline (10.35 ± 2.227, *n*=9)<br>One-way ANOVA with Dunnett's multiple comparisons test<br>$F(3, 31) = 6.839$, $p = 0.0011$<br>Naïve vs. Citrulline, $p = 0.3543$; Naive vs. Aspartate, $p = 0.0324$; Naïve vs. Aspartate + Citrulline, $p = 0.0004$ |
| 5B | WT + SCR shRNA (5.248 ± 0.5980, *n*=64), TG + SCR shRNA (602.4 ± 33.67, *n*=78), TG + ODC1 shRNA (49.56 ± 3.076, *n*=83)<br>One-way ANOVA with Tukey's multiple comparisons test<br>$F(2, 222) = 270.5$, $p < 0.0001$<br>WT + SCR shRNA vs. TG + SCR shRNA, $p < 0.0001$; TG + SCR shRNA vs. TG + ODC1 shRNA, $p < 0.0001$ |
| 5D | WT + SCR shRNA (499701 ± 31777, *n*=66), TG + SCR shRNA (2269166 ± 154816, *n*=77), TG + ARG1 shRNA (541016 ± 34906, *n*=63), TG + ODC1 shRNA (920630 ± 63797, *n*=42)<br>One-way ANOVA with Tukey's multiple comparisons test<br>$F(3, 244) = 77.94$, $p < 0.0001$<br>WT + SCR shRNA vs. TG + SCR shRNA, $p < 0.0001$; TG + SCR shRNA vs. TG + ARG1 shRNA, $p < 0.0001$; TG + SCR shRNA vs. TG + ODC1 shRNA, $p < 0.0001$ |
| 5E | WT + SCR shRNA (1083027 ± 67538, *n*=70), TG + SCR shRNA (3477401 ± 230971, *n*=86), TG + ARG1 shRNA (1064900 ± 99093, *n*=63), TG + ODC1 shRNA (1335742 ± 98261, *n*=70)<br>One-way ANOVA with Dunnett's multiple comparisons test<br>$F(3, 285) =62.37$ $p < 0.0001$<br>WT + SCR shRNA vs. TG + SCR shRNA, $p < 0.0001$; TG + SCR shRNA vs. TG + ARG1 shRNA, $p < 0.0001$; TG + SCR shRNA vs. TG + ODC1 shRNA, $p < 0.0001$ |

| | |
|---|---|
| 5H | WT + SCR shRNA (18.88 ± 1.231, $n$=16), TG + SCR shRNA (28.66 ±1.267, $n$=15), TG + ARG1 shRNA (20.73 ± 1.123, $n$=14), TG + ODC1 shRNA (22.65 ± 1.459, $n$=19)<br>One-way ANOVA with Tukey's multiple comparisons test<br>$F(3, 60) = 9.912$, $p < 0.0001$<br>WT + SCR shRNA vs. TG + SCR shRNA, $p < 0.0001$; TG + SCR shRNA vs. TG + ARG1 shRNA, $p = 0.0008$; TG + SCR shRNA vs. TG + ODC1 shRNA, $p = 0.0083$ |
| 5I | WT + SCR shRNA (61.31 ± 8.8, $n$=16), TG + SCR shRNA (121.6 ± 8.077, $n$=15), TG + ARG1 shRNA (59.77 ± 5.324, $n$=13), TG + ODC1 shRNA (63.68 ± 6.485, $n$=19)<br>One-way ANOVA with Tukey's multiple comparisons test<br>$F(3, 59) = 15.67$, $p < 0.0001$<br>WT + SCR shRNA vs. TG + SCR shRNA, $p < 0.0001$; TG + SCR shRNA vs. TG + ARG1 shRNA, $p < 0.0001$; TG + SCR shRNA vs. TG + ODC1 shRNA, $p < 0.0001$ |
| 5K | WT + SCR shRNA (46.08 ± 0.4026, $n$=178), TG + SCR shRNA (56.51 ± 0.4697, $n$=217), TG + ARG1 shRNA (39.22 ± 0.5811, $n$=158), TG + ODC1 shRNA (42.49 ± 0.7647, $n$=130)<br>One-way ANOVA with Tukey's multiple comparisons test<br>$F(3, 679) = 215.1$, $p < 0.0001$<br>WT + SCR shRNA vs. TG + SCR shRNA, $p < 0.0001$; TG + SCR shRNA vs. TG + ARG1 shRNA, $p < 0.0001$; TG + SCR shRNA vs. TG + ODC1 shRNA, $p < 0.0001$ |
| 5L | WT + SCR shRNA (70.87 ± 1.326, $n$=180), TG + SCR shRNA (89.77 ± 1.042, $n$=216), TG + ARG1 shRNA (71.45 ± 1.075, $n$=158), TG + ODC1 shRNA (83.67 ± 1.046, $n$=129)<br>One-way ANOVA with Tukey's multiple comparisons test<br>$F(3, 679) = 71.61$, $p < 0.0001$<br>WT + SCR shRNA vs. TG + SCR shRNA, $p < 0.0001$; TG + SCR shRNA vs. TG + ARG1 shRNA, $p < 0.0001$; TG + SCR shRNA vs. TG + ODC1 shRNA, $p = 0.0017$ |
| 6B | **Tonic current**:<br>Control (10.37 ± 2.441, $n$=11), ABH (2.109 ± 0.3597, $n$=5), DFMO (3.655 ± 0.7894, $n$=10)<br>One-way ANOVA with Tukey's multiple comparisons test<br>$F(2, 23) = 5.463$, $p = 0.0114$<br>Control vs. ABH, $p = 0.0297$; Control vs. DFMO, $p = 0.0292$ |
| | **sIPSC amplitude**:<br>Control (41.21 ± 5.367, $n$=11), ABH (32.66 ± 2.188, $n$=5), DFMO (49.68 ± 5.183, $n$=10)<br>One-way ANOVA with Tukey's multiple comparisons test<br>$F(2, 23) = 1.944$, $p = 0.01659$<br>Control vs. ABH, $p = 0.5959$; Control vs. DFMO, $p = 0.4653$ |
| | **sIPSC frequency**:<br>Control (2.293 ± 0.3681, $n$=11), ABH (1.580 ± 0.5373, $n$=5), DFMO (2.329 ± 0.3519, $n$=10)<br>One-way ANOVA with Tukey's multiple comparisons test<br>$F(2, 23) = 0.7804$, $p = 0.4700$<br>Control vs. ABH, $p = 0.5092$; Control vs. DFMO, $p = 0.9974$ |
| 6D | **Tonic current**:<br>WT + SCR shRNA (4.001± 0.4549, $n$=15), TG + SCR shRNA (11.24 ± 1.731, $n$=16), TG + ARG1 shRNA (5.091 ± 1.128, $n$=13), TG + ODC1 shRNA (4.075 ± 0.7321, $n$=19)<br>One-way ANOVA with Tukey's multiple comparisons test<br>$F(3, 59) = 9.969$, $p < 0.0001$<br>WT + SCR shRNA vs. TG + SCR shRNA, $p = 0.0001$; TG + SCR shRNA vs. TG + ARG1 shRNA, $p = 0.0023$; TG + SCR shRNA vs. TG + ODC1 shRNA, $p < 0.0001$ |
| | **sIPSC amplitude**:<br>WT + SCR shRNA (34.57± 1.908, $n$=10), TG + SCR shRNA (41.39 ± 4.835, $n$=16), TG + ARG1 shRNA (37.60 ± 2.878, $n$=12), TG + ODC1 shRNA (38.34 ± 2.662, $n$=17)<br>One-way ANOVA with Tukey's multiple comparisons test<br>$F(3, 51) = 0.5640$, $p = 0.6412$ |

| | |
|---|---|
| | WT + SCR shRNA vs. TG + SCR shRNA, *p* = 0.5821; TG + SCR shRNA vs. TG + ARG1 shRNA, *p* = 0.8771; TG + SCR shRNA vs. TG + ODC1 shRNA, *p* = 0.9116 |
| | **sIPSC frequency**:<br>WT + SCR shRNA (1.983 ± 0.4121, *n*=11), TG + SCR shRNA (2.428 ± 0.3485, *n*=15), TG + ARG1 shRNA (2.058 ± 0.4730, *n*=10), TG + ODC1 shRNA (1.517 ± 0.1954, *n*=15)<br>One-way ANOVA with Tukey's multiple comparisons test<br>F(3, 47) = 1.358, *p* = 0.2670<br>WT + SCR shRNA vs. TG + SCR shRNA, *p* = 0.8036; TG + SCR shRNA vs. TG + ARG1 shRNA, *p* = 0.8848; TG + SCR shRNA vs. TG + ODC1 shRNA, *p* = 0.1991 |
| 6G | WT + SCR shRNA (0.7889 ± 0.1111, *n*=9), TG + SCR shRNA (0.2857 ± 0.1844, *n*=7), TG + ARG1 shRNA (0.9100 ± 0.0481, *n*=10), TG + ODC1 shRNA (0.9364 ± 0.0543, *n*=11)<br>One-way ANOVA with Tukey's multiple comparisons test<br>F(3, 33) = 8.275, *p* = 0.0003<br>WT + SCR shRNA vs. TG + SCR shRNA, *p* = 0.0093; TG + SCR shRNA vs. TG + ARG1 shRNA, *p* = 0.0008; TG + SCR shRNA vs. TG + ODC1 shRNA, *p* = 0.0004 |
| 6I | **% Alteration**:<br>WT + SCR shRNA (62.27 ± 4.778, N=9), TG + SCR shRNA (39.43 ± 3.669, N=7), TG + ARG1 shRNA (62.05 ± 3.468, N=9), TG + ODC1 shRNA (61.02 ± 4.770, N=9)<br>One-way ANOVA with Tukey's multiple comparisons test<br>F(3, 30) = 5.910, *p* = 0.0027<br>WT + SCR shRNA vs. TG + SCR shRNA, *p* = 0.0057; TG + SCR shRNA vs. TG + ARG1 shRNA, *p* = 0.0062; TG + SCR shRNA vs. TG + ODC1 shRNA, *p* = 0.0094 |
| | **Number of entries**:<br>WT + SCR shRNA (20.78 ± 1.267, N=9), TG + SCR shRNA (22.43 ± 1.645, N=7), TG + ARG1 shRNA (20.22 ± 1.115, N=9), TG + ODC1 shRNA (18.33 ± 1.500, N=9)<br>One-way ANOVA with Tukey's multiple comparisons test<br>F(3, 30) = 1.427, *p* = 0.2543<br>WT + SCR shRNA vs. TG + SCR shRNA, *p* = 0.8454; TG + SCR shRNA vs. TG + ARG1 shRNA, *p* = 0.6961; TG + SCR shRNA vs. TG + ODC1 shRNA, *p* = 0.1998 |
| 6J | **Day 1. Acquisition**:<br>WT + SCR shRNA (30.01 ± 4.152, N=11), TG + SCR shRNA (46.08 ± 9.101, N=10), TG + ARG1 shRNA (79.42 ± 19.67, N=10), TG + ODC1 shRNA (66.44 ± 16.34, N=10)<br><br>**Day2. Retrieval**<br>WT + SCR shRNA (441.3 ± 48.84, N=11), TG + SCR shRNA (106.5 ± 14.03, N=10), TG + ARG1 shRNA (303.7 ± 46.59, N=10), TG + ODC1 shRNA (311.8 ± 46.18, N=10)<br>One-way ANOVA with Tukey's multiple comparisons test<br>F(3, 37) = 10.98, *p* < 0.0001<br>WT + SCR shRNA vs. TG + SCR shRNA, *p* < 0.0001; TG + SCR shRNA vs. TG + ARG1 shRNA, *p* = 0.0115; TG + SCR shRNA vs. TG + ODC1 shRNA, *p* = 0.0081 |
| 7C | WT + SCR shRNA (0.7778 ± 0.4339, *n*=9), TG + SCR shRNA (32.64 ± 3.095, *n*=14), TG + ARG1 shRNA (24.36 ± 2.667, *n*=11), TG + ODC1 shRNA (18.73 ± 1.442, *n*=15)<br>One-way ANOVA with Tukey's multiple comparisons test<br>F(3, 45) = 29.10, *p* < 0.0001<br>WT + SCR shRNA vs. TG + SCR shRNA, *p* < 0.0001; TG + SCR shRNA vs. TG + ARG1 shRNA, *p* = 0.0692; TG + SCR shRNA vs. TG + ODC1 shRNA, *p* = 0.0002 |

| | |
|---|---|
| S2B | **Citrulline**:<br>Control (0.09098 ± 0.009592, *n*=5), $^{15}NH_4Cl$ (0.1 ± 0.01477, *n*=4)<br>Unpaired two-tailed t-test, $p = 0.6064$ |
| | **Arginine**:<br>Control (0.1007 ± 0.0008772, *n*=5), $^{15}NH_4Cl$ (0.09983 ± 0.001328, *n*=4)<br>Unpaired two-tailed t-test, $p = 0.5944$ |
| | **Ornithine**<br>Control (0.1600 ± 0.001789, *n*=5), $^{15}NH_4Cl$ (0.1933 ± 0.008056, *n*=4)<br>Unpaired two-tailed t-test, $p = 0.0027$ |
| | **Putrescine**<br>Control (0.08814 ± 0.001434, *n*=5), $^{15}NH_4Cl$ (0.1088 ± 0.00366, *n*=4)<br>Unpaired two-tailed t-test, $p = 0.0007$ |
| | **GABA**:<br>Control (0.1126 ± 0.001939, *n*=5), $^{15}NH_4Cl$ (0.2813 ± 0.006588, *n*=4)<br>Unpaired two-tailed t-test, $p < 0.0001$ |
| S3C | Aβ (7.935 ± 1.667, n=16), Aβ + CQ (2.077 ± 0.6188, n=6)<br>Unpaired two-tailed t-test, $p = 0.0485$ |
| S3D | Aβ (100 ± 5.264, *n*=6), Aβ + DFMO (88.79 ± 5.497, *n*=6), Aβ + ABH (94.13 ± 4.511, *n*=6), Aβ + KDS2010 (87.42 ± 3.012, *n*=6),<br>Ordinary two-way ANOVA with Dunnett's multiple comparisons test<br>$F(3, 15) = 4.2$<br>Aβ vs Aβ + DFMO, $p = 0.0323$; Aβ vs. Aβ + ABH, $p = 0.3457$; Aβ vs Aβ + KDS2010 $p = 0.0163$. |
